# Improved post-stroke spontaneous recovery by astrocytic extracellular vesicles

**DOI:** 10.1101/2021.04.19.440321

**Authors:** Yessica Heras-Romero, Axayacatl Morales-Guadarrama, Ricardo Santana-Martínez, Isaac Ponce, Ruth Rincón-Heredia, Augusto César Poot-Hernández, Araceli Martínez-Moreno, Esteban Urrieta, Berenice N. Bernal-Vicente, Aura N. Campero-Romero, Perla Moreno-Castilla, Nigel H. Greig, Martha L. Escobar, Luis Concha, Luis B. Tovar-y-Romo

## Abstract

Spontaneous recovery after a stroke accounts for a major part of the neurological recovery in patients. However limited, the spontaneous recovery is mechanistically driven by axonal restorative processes for which several molecular cues have been previously described. We report the acceleration of spontaneous recovery in a preclinical model of ischemia/reperfusion in rats via a single intracerebroventricular administration of extracellular vesicles released from primary cortical astrocytes. We used MRI, confocal and multiphoton microscopy to correlate the structural remodeling of the corpus callosum and striatocortical circuits with neurological performance over 21 days. We also evaluated the functionality of the corpus callosum by repetitive recordings of compound action potentials to show that the recovery facilitated by astrocytic extracellular vesicles was both anatomical and functional. Our data provide compelling evidence that astrocytes can hasten the basal recovery that naturally occurs post-stroke through the release of cellular mediators contained in extracellular vesicles.

## Introduction

The epidemiology of stroke has dynamically changed over recent decades, during which a constant improvement in clinical care and increased preventive measures, acute treatment, and timely neurorehabilitation have reduced the fatality rate and transformed into a prevailing cause of chronically disabling disease across the developed world ^1^ and several developing nations, particularly in Latin America ^2,3^. Acute ischemic stroke impacts cognition, sensation, vision, language, and motor performance, with prominent signs of hemianopsia (loss of sight in half of the visual field), diplopia (double vision of the same object), speech deficits, paresis (muscular weakness), paresthesia (abnormal sensations of the skin) and other motor and sensory deficits ^4^.

Despite impactful neurological impairment, most patients regain some of their lost neurological functions without intervention via a phenomenon of spontaneous recovery, which is determined exclusively by the passage of time ^5^. Such spontaneous recovery occurs rapidly, at the level of impairment, and is driven by plasticity mechanisms initiated by the stroke ^6^. Functional motor gains related to spontaneous recovery are considered to be due to increased connectivity of motor network areas that were initially disjointed by the ischemic attack ^7^. However, the extent of recovery varies among patients, and the level of neurological regain is considered to be proportional to the initial deficit ^8^. Rehabilitation during a narrow time window can enhance functional recovery, primarily through compensation. However, most clinical trials have failed to show differences in outcome endpoints between experimental and control interventions ^9^.

Studies in animal models attempted to elucidate a time-limited period of heightened plasticity after focal brain injury and the mechanisms behind spontaneous recovery ^6,10^. Cumulative data indicate that axonal repair, neurogenesis, and inflammation resolution mechanistically intervene in this recovery ^11,12^. Particularly, axonal remodeling is a principal mechanism in which surviving neurons in the peri-infarct cortex establish new connections within the motor, somatosensory, and premotor areas within the ipsilesional hemisphere, with some reaching contralesional sites ^13,14^, while others even innervate the frontal motor regions to the brainstem or spinal cord ^15^. However, these new projections alter the topography of cortical projections in the somatosensory system and change the aggregate map of projections ^16,17^.

Astrocytes and astrocytic gliosis are known to impede the recovery of injured tissue by blocking axonal sprouting through the production of structural astrocytic proteins such as glial fibrillary acidic protein (GFAP) and vimentin ^18^, as well as neurite-inhibiting signaling molecules such as Nogo-A ^19^. Nonetheless, astrocytes have multiple essential support functions, whose loss can precipitate or contribute to neurodegeneration ^20^. Astrocytes are more resilient against ischemia than neurons; ^21,22^ therefore, astrocyte survival holds the potential to restore neuronal integrity and promote a functional improvement, especially in the ischemic penumbra ^23^.

Here we evaluated in a preclinical stroke model caused by the transient occlusion of the middle cerebral artery (MCAO) in rat whether astrocytes might influence spontaneous recovery through signaling mechanisms mediated by the release of extracellular vesicles (EVs), which are known to contribute to the modulation of CNS physiology and pathology ^24^.

## Results

### EV release from astrocytes changes after hypoxia

We isolated EVs released from primary astrocytes cultured under normoxic conditions to characterize their effects on stroke evolution. We also subjected astrocytes to a hypoxic challenge to assess whether EVs would retain their molecular activities in response to ischemia. We first ran a physical and molecular characterization of the EVs released from cultured primary cortical astrocytes obtained from P2 newborn rats. The isolation method employed in this study, which involves differential centrifugation, produces a homogeneous EV population with a size consistent with exosomes, as analyzed by transmission electron microscopy (Figure 1a) and nanoparticle tracking analysis (Figure 1c, and d). The isolated EVs also expressed the canonical exosomal marker CD63 (Figure 1b). When we subjected the astrocytic cultures to hypoxia for six h, the number of exosomes collected in 48 h-conditioned media was significantly lower than the normoxic conditions (Figure 1 c, d, and e), so that a physiological outcome to ischemic stress would be reducing the release of EVs.

**Figure 1.**
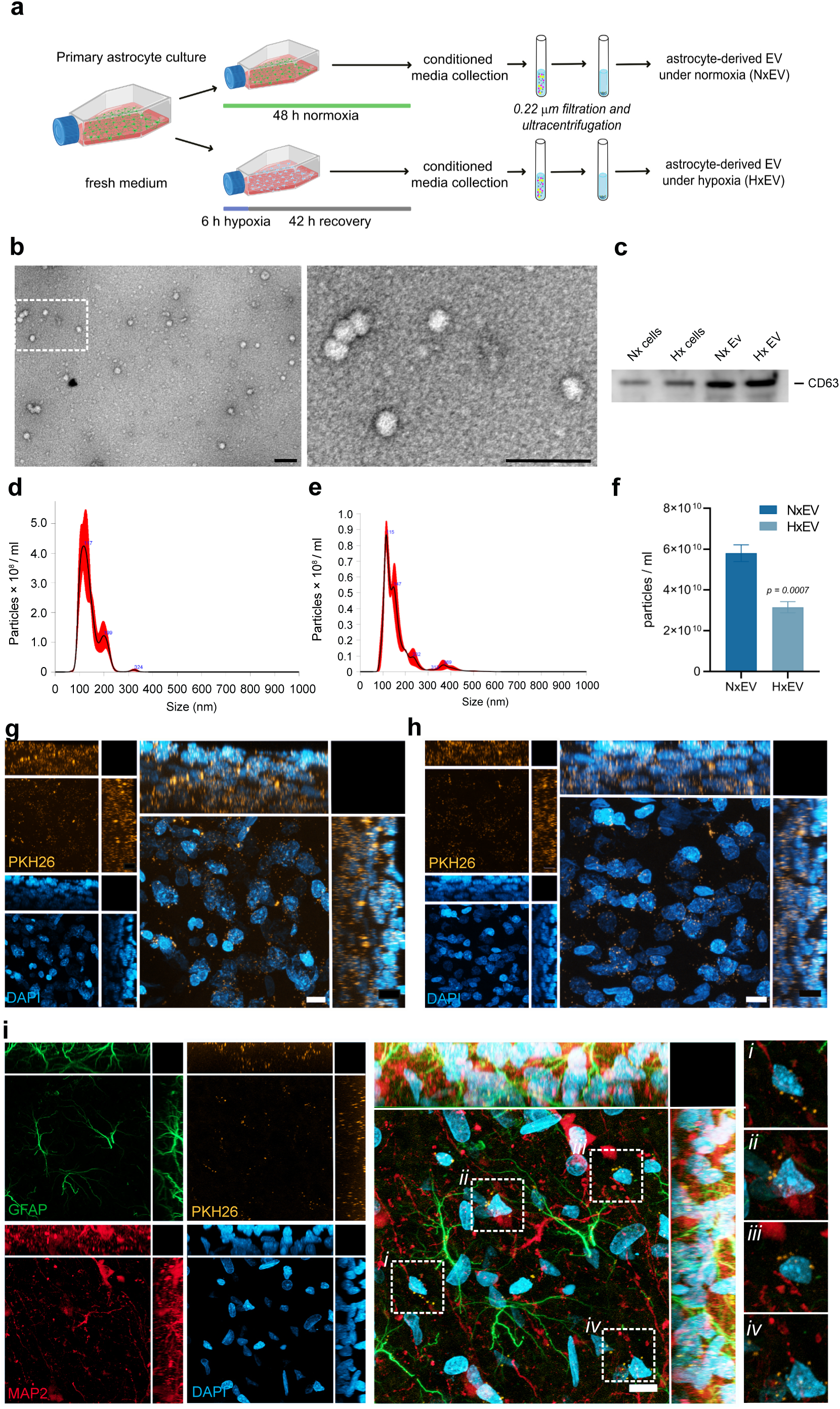
Characterization of exosomes produced by astrocytes. **(a)** Experimental design for EVs collection. **(b)** Transmission electron microscopy micrographs of EVs isolated from astrocyte cultures for 24 h. Vesicles were visualized by negative staining with uranyl formate on copper/carbon-coated grids; scale bars equal 100 nm. **(c)** Immunoblot showing the exosome canonical marker CD63 in protein lysates from EVs and astrocytes cultured under normoxia (Nx) and exposed to 6 h hypoxia and 42 h of recovery (Hx). Size distribution of particles in suspensions by nanoparticle tracking analysis of EVs isolated under normoxia for 48 h **(d)** and after a 6 h hypoxic stimulus and 42 h recovery **(e)**. The graph in **(f)** shows the difference in the number of EVs released from astrocytes subjected to hypoxia (HxEV) and controls (NxEV) for 48 h. Data represent the mean ± standard deviation of three independent measurements. Distribution of EVs stained with PKH26 (orange) injected i.c.v. into the brain of rats within the striatum **(g)** and motor cortex **(h)**. **(i)** EVs internalized in neurons (MAP2; red) and astrocytes (GFAP; green) and preferentially localized to perinuclear (DAPI; blue) regions. Dotted squares demark regions where EVs localize, images on the left are magnifications of those regions. Images in **(g, h, and i)** are maximum projections of a Z-stack of 20 optical slices showing the orthogonal planes, and nuclei are stained with DAPI (blue), scale bar equals 10 µm.

### Administration of astrocyte-derived EVs in the rat brain

In a pilot experiment, using a pulled glass microcapillary pipette, we administered stereotaxic injections of EVs collected from astrocyte-conditioned media resuspended in 0.1 mM PB into the lateral ventricle of the rat brain. We evaluated EV boluses with total protein concentrations of 200 and 400 ng determined by bicinchoninic acid protein assay in a volume of 4 µl. The higher concentration (400 ng) yielded a larger difference in infarct volume respective to vehicle-injected controls and was therefore used for the rest of the study. On average, we injected approximately 8.5×10^7^ vesicles in each administration. Subsequently, we explored the distribution of EVs in the rat brain. We noted that exogenous EVs reached the striatum (Figure 1f) and motor cortex (Figure 1g) within 2 h (Supplementary Figure 1) and remained identifiable 24 h post-injection. Further, we found that EVs became internalized in neurons and astrocytes (Figure 1h, Supplementary Figure 1) and were preferentially localized in the perinuclear region. Using these experimental parameters, we studied the effects of EVs produced by astrocytes cultured under normoxic conditions for 48 h (NxEV) and those released by astrocytes over the same period after culturing under hypoxic conditions for six h (HxEV) to determine whether hypoxia modifies the neuroprotective potential of astrocyte EVs.

### Spontaneous recovery in the preclinical model of stroke

We began by determining the temporal evolution of infarct assessed with MRI. Figure 2a shows coronal slices from a T2-weighted MRI sequence of stroke-challenged animals at 24 h with NxEV or HxEV administration compared to control animals injected with vehicle alone. Evident at 24 h, there is a significant reduction in the average infarct volume in animals that received either type of EVs compared to the control group (p = 0.0157 for NxEV and p = 0.0163 for HxEV two-way ANOVA with Tukey, Figure 2a and 2c). At this time, there was no difference between NxEV and HxEV (p=0.9997), but a difference became apparent over time. Specifically, by day 21 after stroke, p values were 0.0144 for NxEV and <0.0001 for HxEV compared to the control and 0.002 between NxEV and HxEV (Figure 2b and 2c), demonstrating that a single injection of EVs at the beginning of the reperfusion phase contributed to the gradual reduction of the infarct volume. These observations correlated with the natural occurrence of spontaneous recovery in the rat after MCAO. We assessed the neurological performance of subjects on a series of tests that assess the integrity of sensory, motor, and sensorimotor circuits (Table 1). Control stroke-challenged rats that received vehicle alone demonstrated a significant recovery (p=0.0374, t=3.576, df=3 in paired t-test) over 21 d in the neurological parameters examined, as compared to the values obtained by each rat at 24 h after stroke (Figure 2d, 2e and Supplementary Figure 2). Neurological recovery was enhanced in the groups that received EVs; NxEV p=0.0046, t=7.667, df=3, and HxEV p=0.0032, t=8.66, df=3 (Figure 2d, 2e, and Supplementary Figure 2).

**Figure 2.**
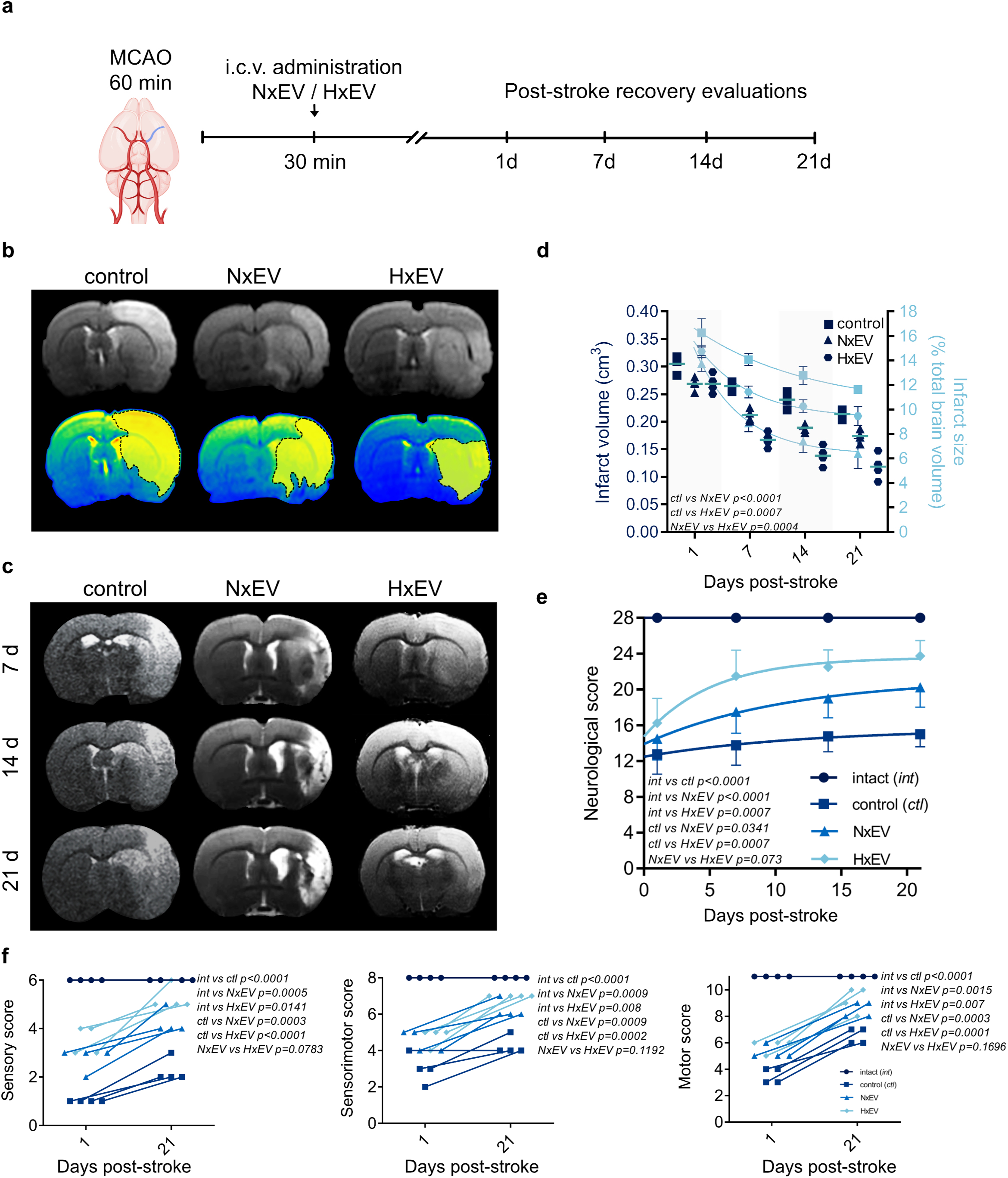
Intracerebroventricular administration of EVs shed by cultured astrocytes reduces the infarct volume in rats subjected to MCAO. **(a)** Time frame of the experimental design. **(b)** Determination of infarct volume by magnetic resonance imaging (MRI) 24 h after MCAO in rats treated with vehicle (control), EVs derived from astrocytes grown under normoxic (NxEV) and hypoxic (HxEV) conditions. The figure shows on top the coronal section of T2 weighted sequences at AP -0.5 from Bregma, and at the bottom, the infarcted area is shown in a yellow mask over a converted image that shows the regions of hyperdensity in a color scale. **(c)** Time course of MRI images of the different groups at 7, 14, and 21 days post-stroke. MRI shows the affected striatum and adjacent premotor, primary motor, and somatosensory cortices and reduced affected areas over time. **(d)** Quantification of the infarct volumes determined with whole-brain measurements and the percentage of brain volume affected by stroke. Data show the mean ± SEM of n=4 animals per group. Two-way repeated measures ANOVA followed by Tukey post hoc. **(e)** The figure shows the evolution of neurological performance, and thus the motor and sensory recovery over 21 d of rats in the indicated experimental groups. Data points are the mean ± SEM of 4 rats per group followed over time. Two-way repeated measures ANOVA followed by Tukey post hoc. **(f)** The neurological evaluation was composed of tests that assessed performance in sensory and motor tasks and the integration of both. The figures show the initial (24 h) and last assessments (21 d) of each rat in each evaluation category.

**Table 1.**
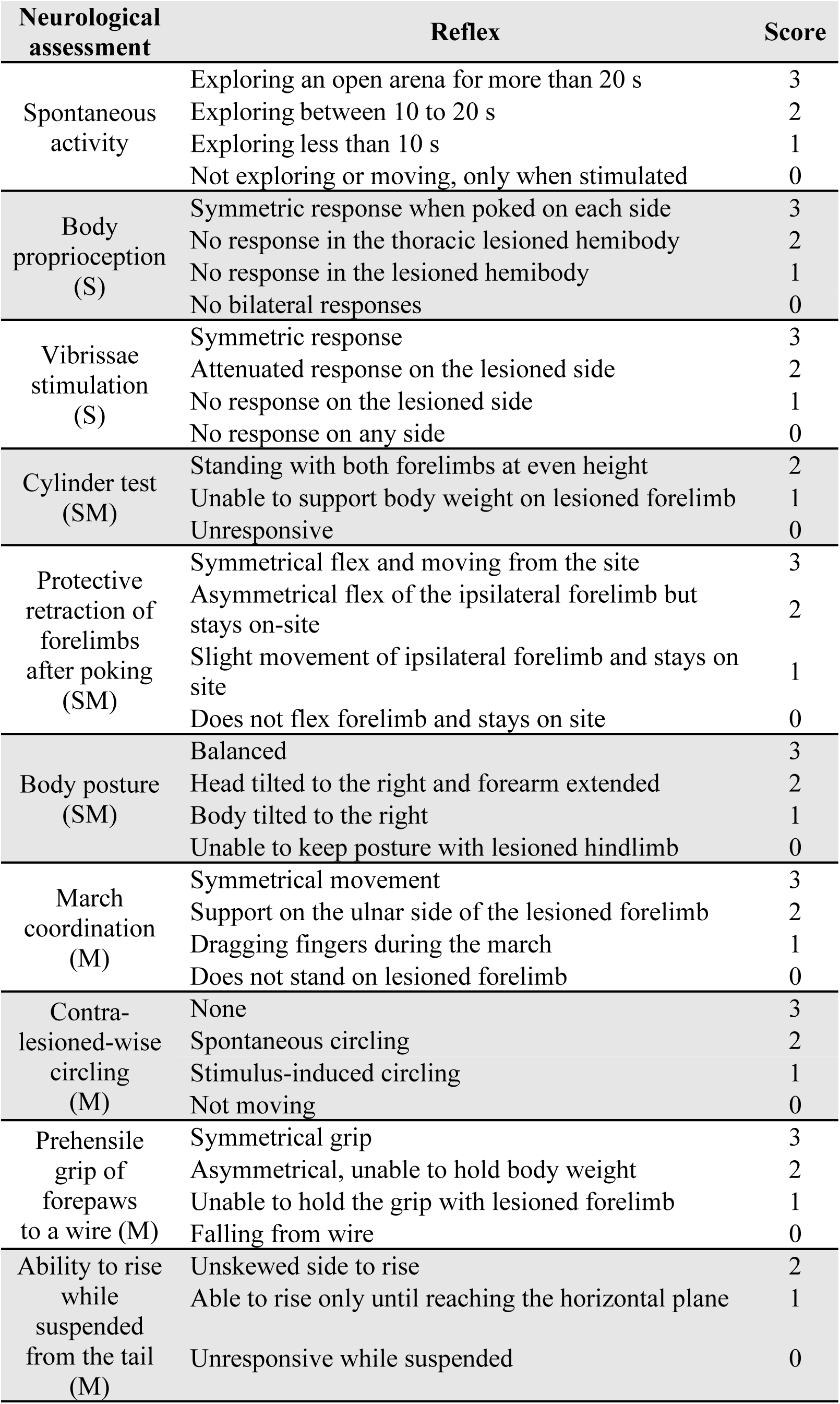
Items evaluated in neurofunctional assessments. Every subject’s neurological performance was assessed at 1, 7, 14, and 21 d after stroke in ten items, and for each one, a corresponding value was assigned. (S) somatosensory evaluation, (M) motor evaluation, (SM) evaluation of the integration of somatosensory/motor coordination. Behavioral evaluations were cross-validated by a trained analyst blind to the experimental conditions.

| Neurological assessment | Reflex | Score |
| --- | --- | --- |
| Spontaneous activity | Exploring an open arena for more than 20 s | 3 |
|  | Exploring between 10 to 20 s | 2 |
|  | Exploring less than 10 s | 1 |
|  | Not exploring or moving, only when stimulated | 0 |
| Body proprioception (S) | Symmetric response when poked on each side | 3 |
|  | No response in the thoracic lesioned hemibody | 2 |
|  | No response in the lesioned hemibody | 1 |
|  | No bilateral responses | 0 |
| Vibrissae stimulation (S) | Symmetric response | 3 |
|  | Attenuated response on the lesioned side | 2 |
|  | No response on the lesioned side | 1 |
|  | No response on any side | 0 |
| Cylinder test (SM) | Standing with both forelimbs at even height | 2 |
|  | Unable to support body weight on lesioned forelimb | 1 |
|  | Unresponsive | 0 |
| Protective retraction of forelimbs after poking (SM) | Symmetrical flex and moving from the site | 3 |
|  | Asymmetrical flex of the ipsilateral forelimb but stays on-site | 2 |
|  | Slight movement of ipsilateral forelimb and stays on site | 1 |
|  | Does not flex forelimb and stays on site | 0 |
| Body posture (SM) | Balanced | 3 |
|  | Head tilted to the right and forearm extended | 2 |
|  | Body tilted to the right | 1 |
|  | Unable to keep posture with lesioned hindlimb | 0 |
| March coordination (M) | Symmetrical movement | 3 |
|  | Support on the ulnar side of the lesioned forelimb | 2 |
|  | Dragging fingers during the march | 1 |
|  | Does not stand on lesioned forelimb | 0 |
| Contra-lesioned-wise circling (M) | None | 3 |
|  | Spontaneous circling | 2 |
|  | Stimulus-induced circling | 1 |
|  | Not moving | 0 |
| Prehensile grip of forepaws to a wire (M) | Symmetrical grip | 3 |
|  | Asymmetrical, unable to hold body weight | 2 |
|  | Unable to hold the grip with lesioned forelimb | 1 |
|  | Falling from wire | 0 |
| Ability to rise while suspended from the tail (M) | Unskewed side to rise | 2 |
|  | Able to rise only until reaching the horizontal plane | 1 |
|  | Unresponsive while suspended | 0 |

### Hastened recovery of structural alterations by the administration of astrocyte-derived EVs

Using MRI, we studied the impact of EVs from cultured astrocytes on the evolution of the brain’s physical recovery from stroke in a longitudinal study. We measured mean diffusivity, which is a quantitative measure that reflects cellular and membrane density—an increase of mean diffusivity results from edema and necrosis. The basal values of mean diffusivity ranged around 6×10^-4^ mm^2^/ms in the corpus callosum, 3 ×10^-4^ mm^2^/ms in the striatum, and 4 ×10^-4^ mm^2^/ms in the cortex. The values almost doubled in stroke-challenged rats at 24 h after the infarction, irrespective of treatment (Figure 3a), across all brain structures analyzed. Notably, these values declined gradually and were decisively lower at 7 d post-stroke in the animals that received EVs. This was particularly evident in the group that received HxEV, which showed baseline mean diffusivity values by day 21. The corpus callosum and striatum exhibited the most robust recovery rates in this parameter, coincidentally with structures with a lower cellular density than the cortex.

**Figure 3.**
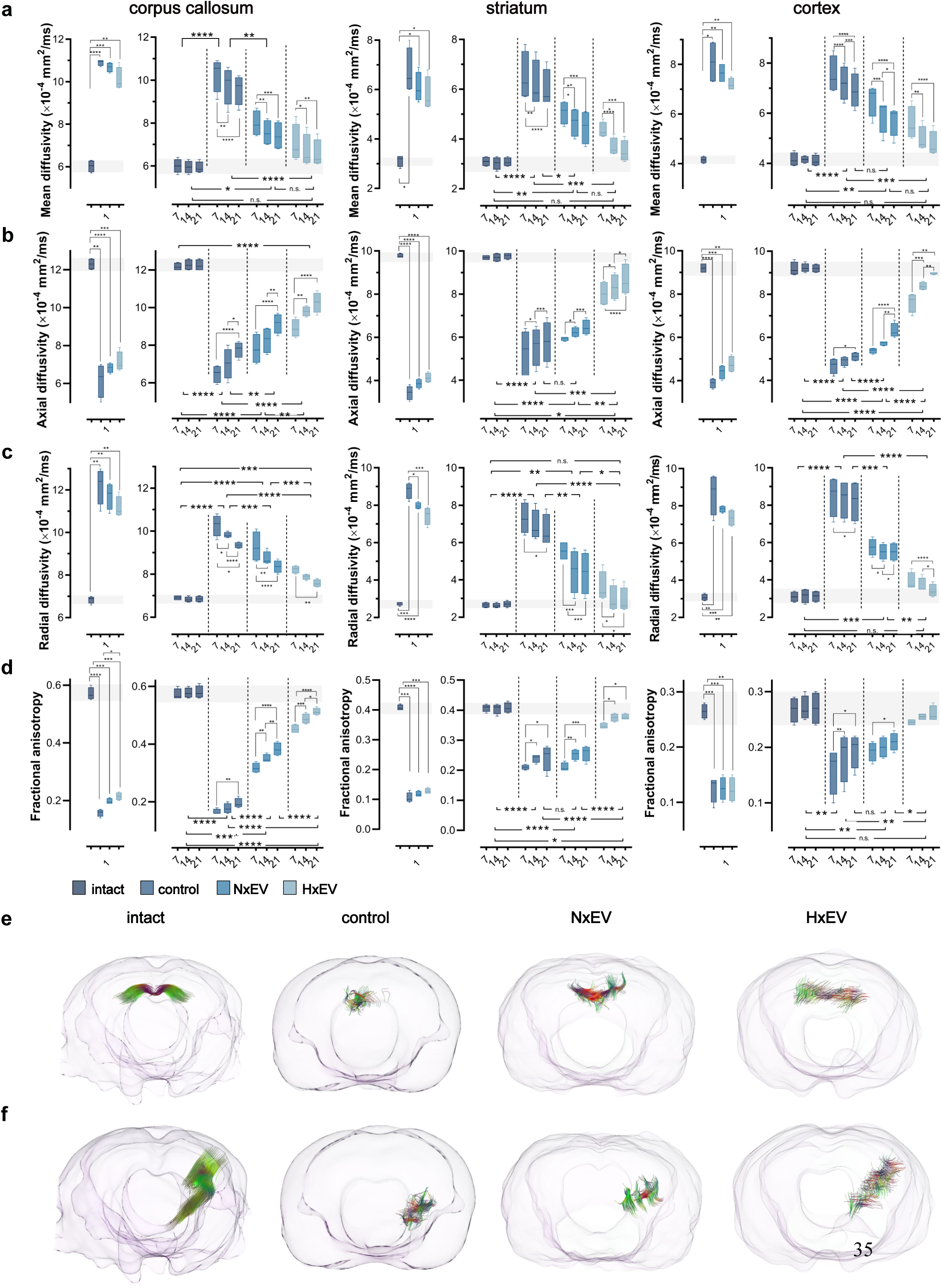
The administration of astrocyte-derived exosomes preserves the structural integrity of neuronal tracts. **(a)** mean diffusivity, **(b)** axial diffusivity, **(c)** radial diffusivity, and **(d)** fractional anisotropy were determined from diffusion tensor imaging (DTI) of the ipsilateral corpus callosum (left column), striatum (middle column), and motor cortex (right column) at 1, 7, 14 and 21 days post-stroke. Boxplots on day 1 show the alterations caused by the stroke in all four DTI parameters; no statistical differences exist between stroke-challenged animals treated with vehicle (control) and those that received EVs 30 min after the beginning of reperfusion. From day 7 onwards, boxplots show the evolution of the recovery hastened by the administration of NxEV or HxEV. Boxplots show the min and max values within each group, the dispersion span from Q1 to Q3, and the mean, n=4. The shaded horizontal bar in each plot marks the span of ± 1 S.D. of the intact group baseline values. Statistical differences of the recovery trend are indicated among groups with two-way repeated measures ANOVA followed by Tukey’s post hoc, and changes over time within each group are also indicated with two-way ANOVA followed by Tukey’s post hoc. * p<0.05, **p<0.01, ***p<0.001 and ****p<0.0001. Fiber tractography produced from DTI of the corpus callosum (e) and a striatocortical tract of the ipsilesional hemisphere (f) of a representative rat of each experimental group 21 d after stroke. Colors in the tracts indicate the orientation of the fiber: transverse fibers (red), anteroposterior fibers (green), and craniocaudal fibers (blue).

As noted above, the underlying processes of spontaneous recovery are firmly bound to axonal damage recovery. Axial diffusivity results from the structural integrity of axons and a decrease of this value mirrors axonal damage. We found that at 24 h post-stroke, all animals displayed a significant reduction in axial diffusivity that was more severe in the striatum>cortex>corpus callosum, which corresponds to the hierarchical structural damage produced by MCAO (Figure 3b). Values in this parameter indicate a time-dependently recovery across all of the groups. However, such recovery was more robust in HxEV, followed by the NxEV group. By day 21 post-stroke, the axial diffusivities of HxEV in the striatum and cortex reached the baseline values; the overall recovery trend of the HxEV group was significantly different from that of the control group (p = 0.0001 for striatum and p < 0.0001 for cortex), while the group with NxEV showed only significant differences in the cortex (p = 0.4259 for striatum and p < 0.0001 for cortex).

Radial diffusivity is another DTI parameter that evaluates axonal integrity, which is increased with demyelination and reduced axonal density. We found this value to be doubled in the corpus callosum, almost quadrupled in the striatum and tripled in the cortex 24 h post-stroke (Figure 3c).

The last parameter that we assessed with DTI was fractional anisotropy (FA), which is related to how freely water molecules move along axons and reflects their structural integrity. A reduction in FA levels indicates axonal damage. Due to the fibrous nature of the corpus callosum, the baseline values of FA in this structure are higher than those of the striatum and the cortex. The FA baseline level in the corpus callosum (about 0.6) drastically dropped to a third in all experimental groups by 24 h post-stroke (Figure 3d). Notably, the administration of HxEV resulted in a substantial but slightly lesser effect than the untreated control group. Whereas FA in the control group slightly recovered by day 21 post-stroke, a very evident recovery occurred with the administration of either NxEV or HxEV. The latter, in particular, demonstrated a more considerable effect and a clear time-dependent trend towards recovery (Figure 3d). The more severely affected striatum exhibited an even more substantial FA decrease 24 h after infarction, and NxEV did not impact the overall recovery trend.

In contrast, HxEV did promote a significant improvement that evolved rapidly at 7 d post-stroke and exhibited almost normal values by day 14. Finally, in the cortex, with a more heterogeneous architecture than the corpus callosum and the striatum, the stroke-induced changes were of lesser magnitude by day 7. HxEV almost fully mitigated them at this early time point (Figure 3d), whereas NxEV did not affect the spontaneous recovery.

Stroke notably impacts the brain as a whole, and structural damage appears not to be restricted to the only sites where blood supply was impeded. Our MRI study shows that the brain hemisphere contralateral to the lesion had structural changes that resolved over time, and the administration of EVs hastened the recovery. EVs induced a full recovery of the mean diffusivity in the corpus callosum by day 21 (Supplementary Figure 3a). The contralateral cortex and striatum showed subtler changes that were normalized by 14 d with HxEV and 21 d with NxEV (Supplementary Figure 3a). Similarly, the axial diffusivities of the corpus callosum and contralateral cortex were fully recovered by day 21 with the administration of EVs (Supplementary Figure 3b). In the striatum, the trend towards full recovery was more marked with HxEV. As in the ipsilateral side, the radial diffusivity was minimally affected by the stroke in the contralateral hemisphere, and the administration of HxEV provided full recovery as early as day seven post-stroke for the striatum and cortex (Supplementary Figure 3c). Finally, the FA of the contralateral structures showed a similar recovery pattern as that of the ipsilateral hemisphere, with the striatum and cortex showing full recovery with the aid of HxEV (Supplementary Figure 3d).

We used vector scalar values from DTI to produce tensor tract reconstructions to provide a graphical visualization of the extent of the recovery of white matter bundles that was facilitated by EVs administration (Supplementary Figure 4 shows the regions of interest where seeds were placed for the tractography). Figure 3e shows DTI-derived tracts across the corpus callosum, and Figure 3f shows a portion of the corticostriatal tract impacted by MCAO. The tractography of intact brains shows a complete reconstruction of the tract segments. In contrast, in the control group, the number of fibers that could be reconstructed was significantly limited, and the shape of reconstructed tracts is highly disorganized and atrophied. In the reconstructions of NxEV and HxEV groups, the length of assembled tracts was longer than that of their respective controls, and the number of fibers was also significantly increased.

### Structural correlations of anatomical fibers with MRI

We evaluated the integrity of brain neuronal structural fibers by immunolabeling microtubule-associated protein 2 (MAP2) and βIII-tubulin (Tuj1). In control animals, there was a near total loss of dendritic staining with MAP2 and all neuronal processes with TUJ1 at 7 d post-stroke (Figure 4). A gradual time-dependent recovery from this loss occurred across all groups, but only to a decidedly limited extent under control conditions. This limit was surpassed with the aid of astrocyte EVs, especially in the cortex (Figure 4). Also illustrated in Figure 4, the recovery of dendritic arborizations was greatly disorganized in the striatum of the control group animals. EVs did not help shape a better organization of the fibers by day 21, as indicated by the directionality of the tracts (Supplementary Figure 5). Overall, the appearance of the neuronal processes labeled with MAP2 and Tuj1 positively correlates with the MRI parameters described above, providing a histological grounding to the determinations made by indirect measurements of DTI.

**Figure 4.**
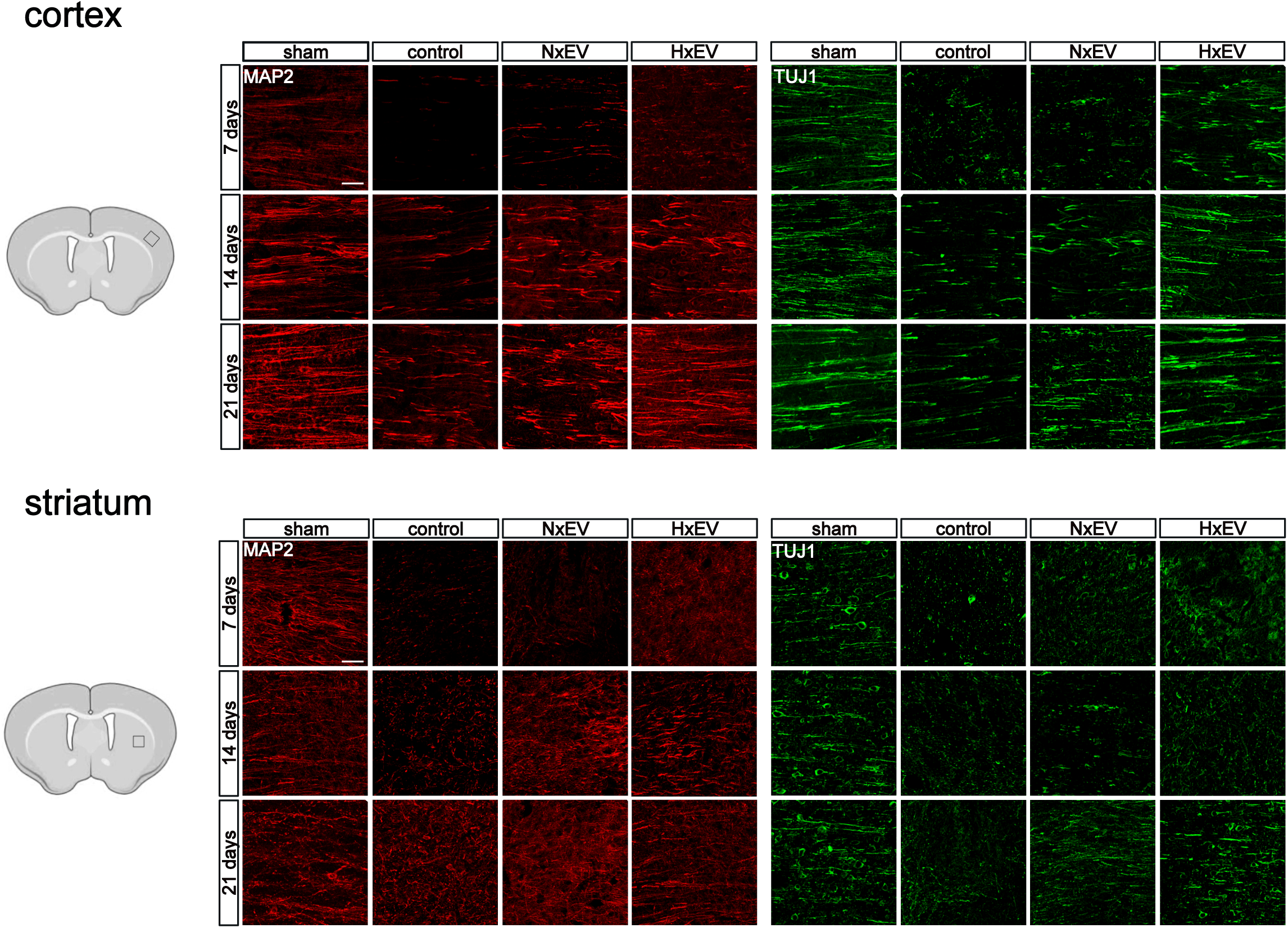
The administration of astrocyte-derived exosomes promotes the recovery of the neuronal processes’ integrity affected by stroke. Representative micrographs of 40 μm-thick immunostained sections with MAP2 (red) and TUJ1 (green) of each experimental group. Images show the evolution of the recovery of the processes in the affected motor cortex and dorsal striatum at 7, 14, and 21 d post-stroke. Microphotographs are Z-stack confocal projections of 10-15 optical slices. Scale bars equal to 50 µm.

Although the administration of EVs facilitated the recovery of the gross neuronal architectural conformation of the brain, we intended to discover whether this treatment also enabled the outgrowth of axons from the lesion core in the striatum towards the innervated cortical areas. We hence at day 14 post-stroke injected the cholera toxin subunit B (Supplementary Figure 6), which is retrogradely transported by the axons, and the fluorescent dye Dil (Figure 5), carried in an anterograde and retrograde fashion through the axonal transport system and followed the localization of the fluorescent label in the innervated cortical regions at day 21 post-stroke.

**Figure 5.**
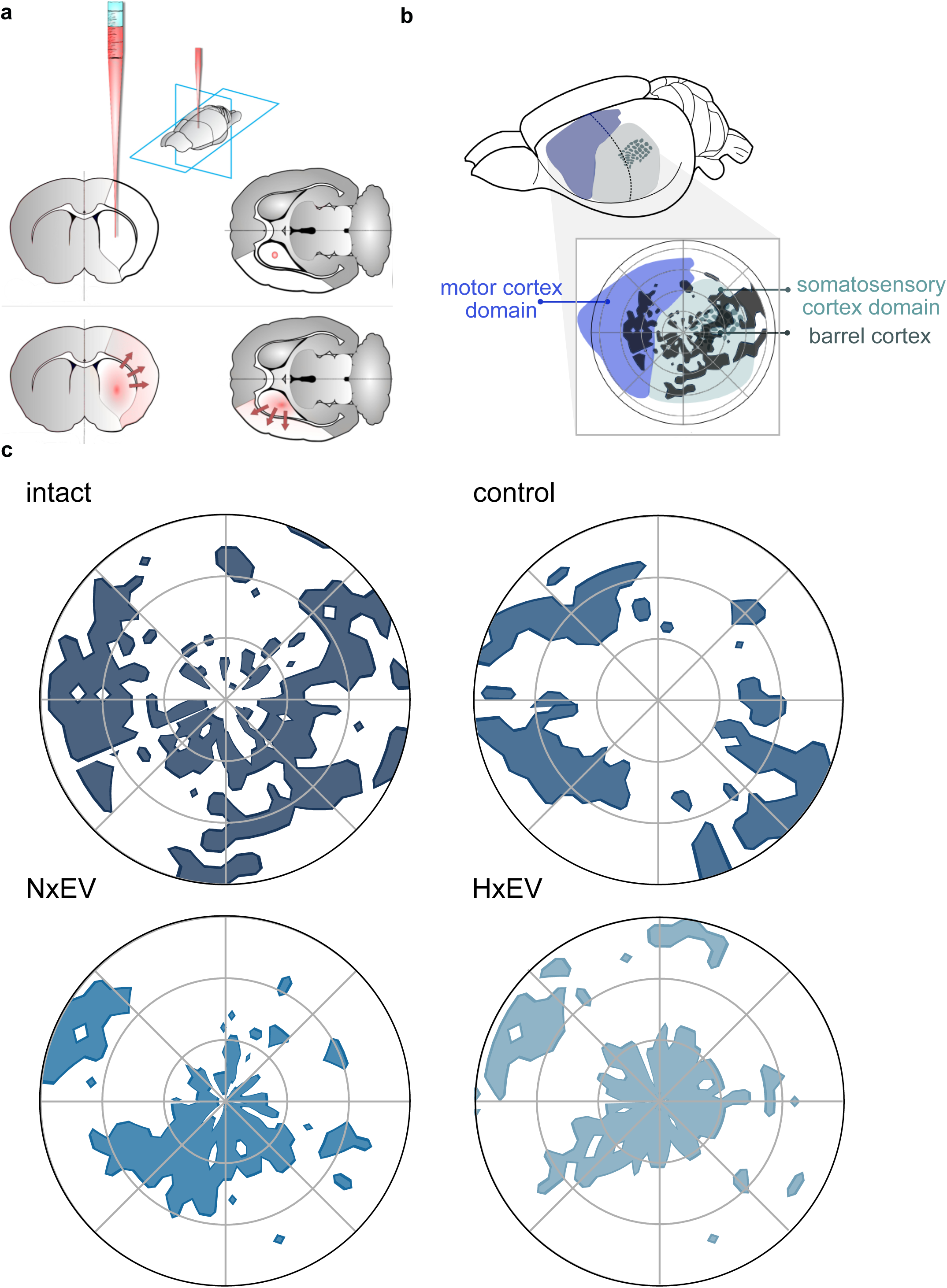
Astrocyte-derived exosomes promote axonal regrowth and the reorganization of cortical innervation maps in the somatosensory cortex. **(a)** Schematic representation of the administration of the fluorescent colorant Dil at 21 d post-stroke and diffusion through axonal transport over 7 d. **(b)** Schematic indication of the localization of polar plots shown in (c) over the cortical territories of motor the motor and somatosensory cortex of the rat, indicated by the localization of the barrel cortex (See Supplementary figure 6). Polar plots of the striatal innervation to the cortex in a representative rat of each group. Each two-photon-captured stack’s maximum projection was converted to the pixels’ cartesian coordinates meeting the threshold set (percentile 90-95 of positive signal). Origin (0, 0) was set the first branch split of the M4 segment in the MCA’s superior trunk. Notice the stroke impact of the somatosensory cortex and the reorganization of the cortical maps produced by EVs.

Next, we determined how the innervation of the ipsilateral cortex from the dorsal striatum was modified by stroke, using two-photon laser scanning confocal microscopy of the entire ipsilateral hemisphere. To achieve this, 20×20 binned images were converted to single pixels and mapped to the cortex in a polar plot with the center located at an anatomical point identifiable by the first major split of the M4 segment of the superior MCA trunk. Overlapping the image of a slice of the corresponding preparation of the contralateral side (corrected for anatomical symmetry) stained with cytochrome C oxidase to distinguish the barrel cortex allowed us to map the general territories of the motor and somatosensory cortices (Supplementary Figure 7). As expected, we found that the stroke caused a drastic loss of striatal projections to the somatosensory and motor cortices. Whereas the administration of EVs did not necessarily rescue the lost innervations, it promoted the reorganization of innervated cortical areas that instead preferentially targeted the parietal somatosensory cortex (Figure 5). The patterns reorganized by EVs were notably similar between normoxia and hypoxia. The motor cortex was predominantly spared in our experiments, possibly owing to the small infarct size on days 14 and 21 (Figure 2).

### Functional axonal regeneration of the corpus callosum

The corpus callosum is the most prominent white matter tract in the mammalian brain, making this structure suitable for the measurement of field potentials in fiber populations. The compound action potentials (CAPs) of the corpus callosum have a two-phase trace with two negative peaks; the first one (N1) representing the myelinated fibers and the second (N2) representing the unmyelinated fibers ^25^ (Figures 6 a and b). The amplitude of CAPs evoked in the corpus callosum reflect the number of contributing axons. The I/O relationship of CAPs evoked in the corpus callosum of the animals showed that EVs administration at 30 min post-injury resulted in significant recovery of CAP amplitude, and this effect was more pronounced in the N2 than N1 CAP component (Figure 6c); thus revealing a greater susceptibility of myelinated fibers to injury. The I/O curves for N1 and N2 amplitudes in the control group were significantly shifted downwards relative to all other groups, indicating an injury-induced reduction of evoked action potentials in the myelinated and unmyelinated axon populations, respectively. In rats from the NxEV and HxEV groups, the N1 and N2 CAP amplitudes were significantly elevated above those of the control group, indicating a favorable degree of neurorestoration of myelinated and unmyelinated fibers. This was more prominent in non-myelinated fibers, revealing a greater susceptibility of myelinated axons.

**Figure 6.**
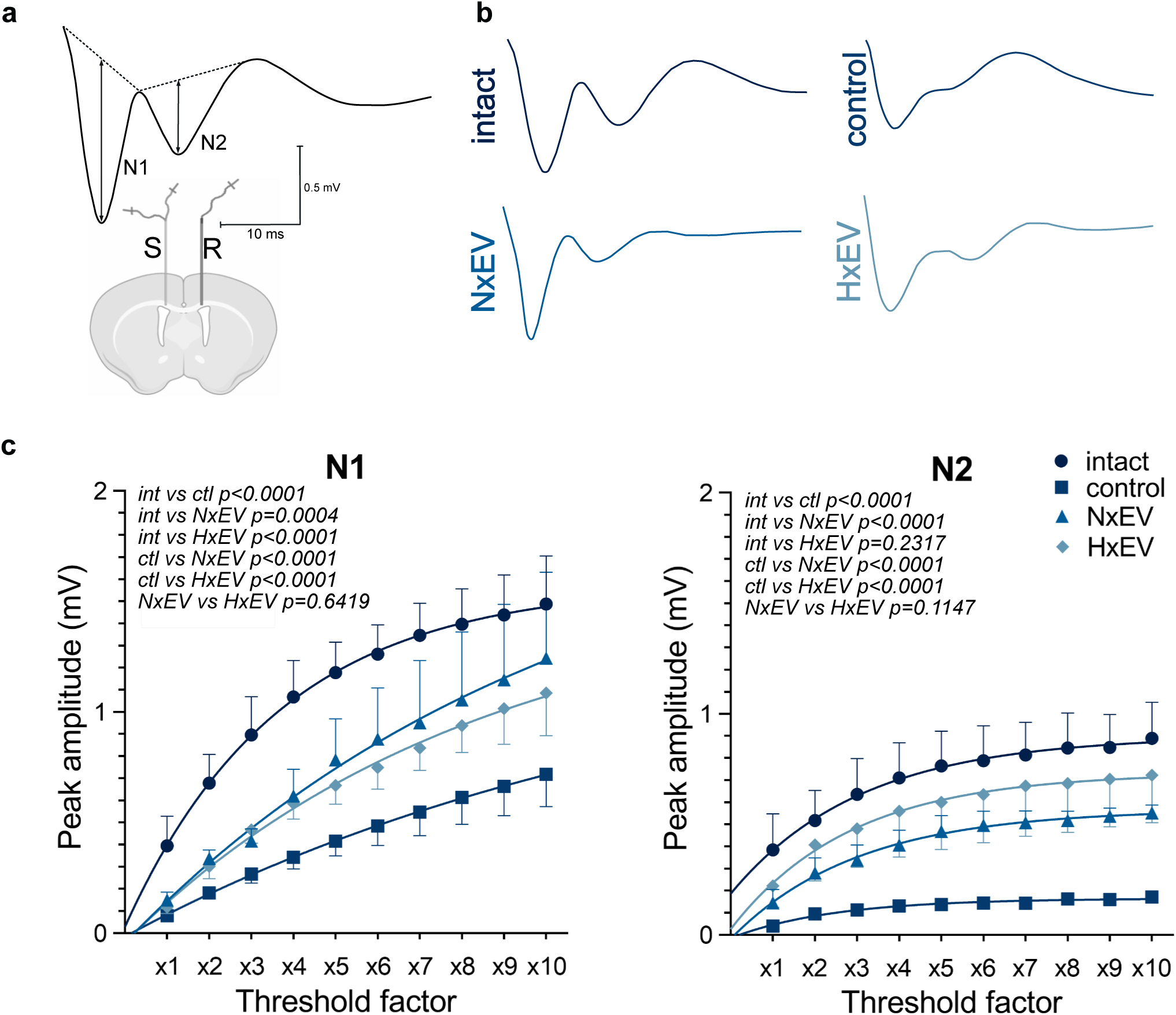
The administration of astrocyte-derived exosomes augments compound action potential recovery of stroke-challenged rats. (a) Schematic representation of electrode placement showing the stimulated (S) and recorded (R) sites in a coronal plane. CAP waveforms display early (N1) and late (N2) negative peaks generated by myelinated and unmyelinated axons, respectively. Dashed lines on CAPs explain the measurements of peak amplitudes from their projected bases. **(b)** Representative CAPs evoked at the maximum stimulus level under each experimental condition. **(c)** Plots for N1 and N2 I/O curves for intact, control, NxEV, and HxEV groups.

The results obtained through the I/O curve measurements reveal a similar recovery pattern to that observed in FA analysis of samples from the corpus callosum (21 d, *i*). Likewise, there was a remarkable correspondence between electrophysiological results (CAPs for N1 at 10X, *ii*) and the functional recovery (combined neurological score at 21 d, *iii*) reported in the present study. Pearson correlation values for *r_i,iii_* = 0.992, p = 0.04, *r_i,ii_* = -0.991, p = 0.043, and *r_i,iii_* = -1, p = 0.003.

### Identification of molecular mediators of axonal remapping within astrocyte-derived EVs cargo

For the last part of this study, we set out to identify the possible molecular mediators of axonal regeneration contained in EVs released from astrocytes. To determine this, we performed a meta-analysis of three protein datasets from previously published astrocyte exosome-derived proteomes with n = 19 ^26^, n = 107 ^27^, and n = 219 ^28^ identified proteins (Figure 7a). We also performed a gene ontology (GO) over-representation test with R (3.6.3) package clusterProfiler ^29^. Subsequently, we selected eight biological processes (BP) GO terms related to axonal growth and synaptogenesis, including synapse organization, response to axon injury, regulation of synapse structure or activity, postsynaptic cytoskeleton organization, postsynapse organization, neuron projection extension, modification of synaptic structure, and axonogenesis (Figure 7b). Unique proteins belonging to these eight BP GO terms (n= 39) were analyzed using STRING with a minimum interaction score of 0.900 and k-means clustering of 3 (Figure 7c).

**Figure 7.**
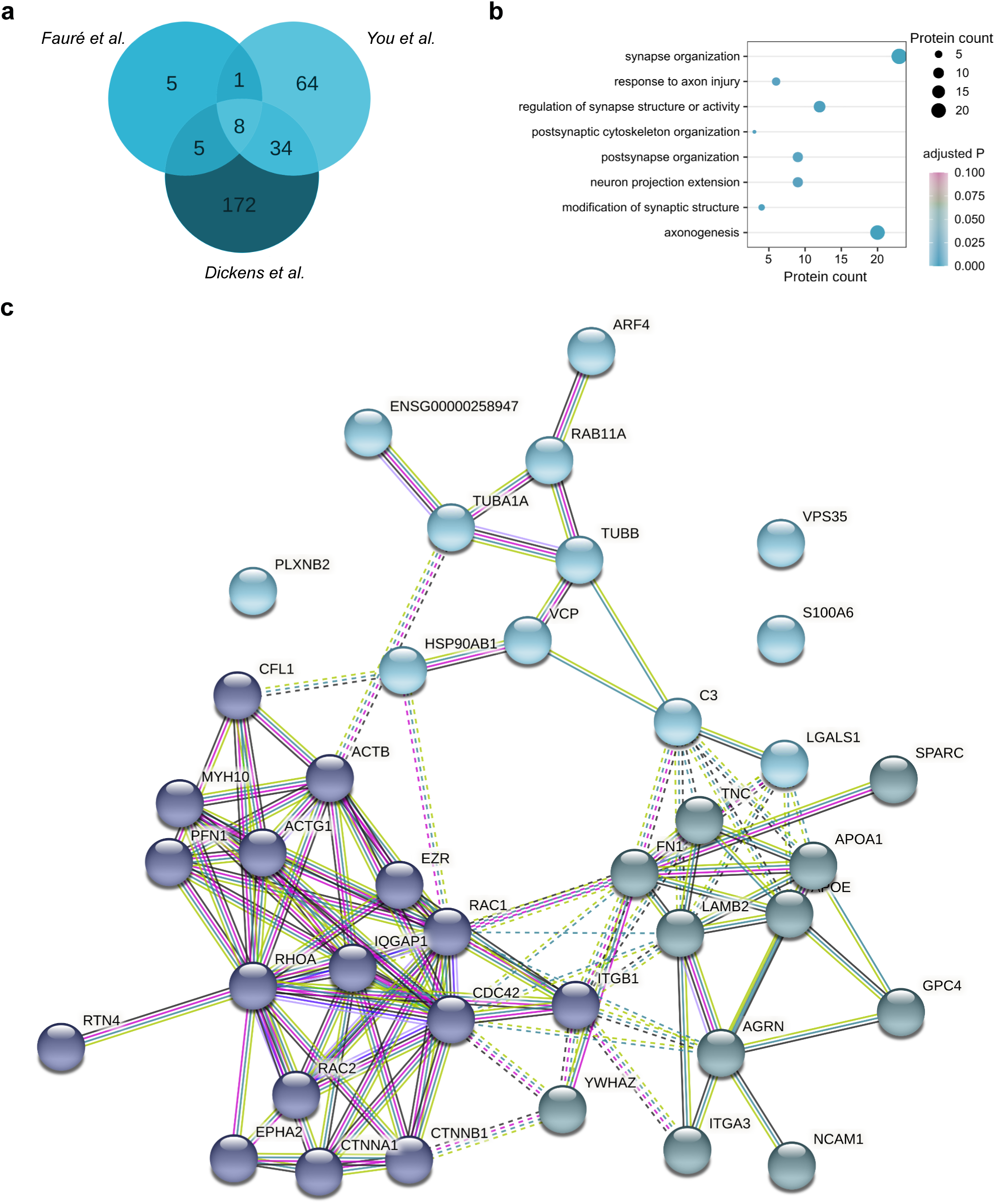
Meta-analysis of astrocytic EV proteomes. **(a)** Venn diagram of the three databases used for the meta-analysis 26–28. **(b)** Biological processes gene ontology analysis of proteins related to neuronal, axonal, and synaptic biological processes. **(c)** STRING analysis of the 39 proteins represented in the BP GO terms in (b), with a minimum interaction score of 0.900 and k-means clustering of 3.

## Discussion

How astrocytes react to ischemic damage after stroke is a complex biological process that involves a coordinated response from a heterogeneous cell population across multiple phases. Here we report that astrocytes produce and release EVs loaded with chemical cues that instigate the axonal remapping of cortical areas damaged by stroke. Importantly, we determined that astrocytes subjected to hypoxic stress release EVs with an increased capacity for repair, although in significantly fewer numbers. Lastly, we found that this mechanism of intercellular communication accelerated the spontaneous recovery of experimental subjects in the subacute phase following stroke. The molecular link between astrocyte-produced biomolecules transmitted in EVs and functional recovery from stroke has not been established before.

The MCAO procedure used in the present study primarily affects motor coordination and stimuli perceptual integration, coded in the circuits of the somatosensory cortex and thalamus ^30^. The subcortical damage affects functional connectivity in the somatosensory cortex, which correlates with cortical activations after electrical stimulation of the affected forelimbs ^31^. Previous work shows that distinct sensorimotor pathways have a significant loss of connectivity two weeks after stroke in the rat ^32^.

During spontaneous recovery, the reorganization of axonal connections and the overall histological patterns of the brain involve either compensation or repair, and the latter is considered to reflect functional recovery ^33^. With work performed on stroke preclinical models, we know that the mechanisms for spontaneous recovery include promoting new brain cortical maps through axon sprouting, remyelination, and blockade of extracellular inhibitory signals ^34,35^. After a stroke, axonal growth and repair mechanisms establish new projections in the contra-lesional hemisphere. The axonal sprouting triggered by stroke generates new local intra-cortical projections as well as long inter-hemispheric projections ^36^, which have been associated with functional recovery ^37^. Through the engagement of both the ipsilateral and contralateral hemispheres, the rostral and caudal portions of the motor cortex are involved in coordinating the skilled reaching performance in the rat ^38^. In humans, the contra-lesional motor cortex plays a central role in the recovery of motor function ^39^, but acute cortical reorganization following focal ischemia appears to occur less rapidly than in rodents ^40^.

Regarding the activators of these repair processes, endogenous cellular and molecular processes occurring during a limited time window promote a regenerative microenvironment in the post-acute ischemic phase. Physiologically, the axon growth cone that covers the terminal zone of neuritic processes is assembled, and along with the reorganization of the cytoskeleton in the proximal axon stump, initiates the regeneration process ^41^. Growth cone formation and axon growth progression are regulated by extracellular factors and intracellular signaling molecules ^42,43^. In addition, it is known that neurons endure profound changes in their transcriptional profile in the subacute phase after stroke, which enables them to undergo plasticity changes ^44^, and astrocytes activate a transcriptional profile that favors the expression of repairing genes ^45^. We hypothesize that astrocytes signal neurons to activate the regenerative mechanisms through intercellular signaling driven by EVs. This type of communication mechanism has recently been shown to be capable of directing these processes. Astrocyte-derived exosomes with prostaglandin D2 synthase expression contribute to axonal outgrowth and functional recovery in stroke by inhibiting the axon growth blocker Semaphorin 3A ^46^, providing evidence that the release of EVs from astrocyte does contribute to axonal growth signaling.

In our meta-analysis of astrocytic EV proteomes, we identified several proteins that regulate axon outgrowth and guidance. In this regard, TUBB is a beta-tubulin protein that is expressed in the developing CNS and is involved in neuronal proliferation, migration, differentiation, and axon guidance ^47^. ACTG1 is a critical protein for axogenesis, axon guidance, and synaptogenesis ^48^. RhoA has been shown to restrict the initiation of neuronal polarization and axon outgrowth during development ^49^, and its knockdown promotes axon regeneration ^50^. RTN4 is a myelin-associated axon growth inhibitor that impairs axon regeneration in the adult mammalian CNS ^51^. Cdc42 targets the cytoskeleton, cell adhesion, and polarity regulators ^52^, and it is involved in axon guidance ^53^. TUBA4A and PFN1 regulate cytoskeletal dynamics ^54^. SPARC is a secreted protein involved in synapse pruning during development ^55^. Rab11A is involved in axon outgrowth ^56^. Ephrin signaling is known to modulate axon guidance and synaptic plasticity and promote long-term potentiation ^57,58^, and its inhibition has been targeted to augment recovery after stroke ^59^.

EVs also regulate several physiological processes by delivering miRNAs to their target cells. Numerous species of these molecules, which were previously found in astrocyte-shed EVs ^28^ are known to regulate axon growth and guidance. miRNAs in distal axons of cortical neurons, like miR15b, miR195, and miR26b, are known to integrate network regulatory systems that promote axonal growth ^60^. Other species that block these processes, like miR203a and miR29a ^60,61^, are decreased in EVs released from astrocytes subjected to proinflammatory signals. Yet other miRNA species, such as miR130a, are known to indirectly regulate VEGFR2 expression ^62^, which is also involved in neuroadaptation that follows stroke ^63^. The axon growth and guidance prompted by EVs most likely result from combining all these different regulatory molecules within the same vesicle. The exact mechanism of action of each miRNA contained within astrocytic EVs requires elucidation.

We enhanced input signals for axonal growth from astrocytes under stress (HxEV) and basal conditions (NxEV), which allowed us, under a permissive environment generated by the stroke, to potentiate the endogenous process of neurite growth and cortical remapping that underlies functional recovery. The endogenous production of EV content within the brain-resident astrocytes changes dynamically throughout recovery after the ischemic insult. By delivering these vesicles 30 min after initiation of the reperfusion phase, we are likely shortening the time taken for the endogenous reparative mechanisms to commence, which is a plausible explanation for the recovery accelerating effect induced by administering astrocytic EVs. The astrocytes in culture were free from the influence of classically activated microglia, which is known to induce A1 reactive astrocytes that promote the death of neurons and oligodendrocytes ^20^.

One of the most important findings of the present study is that astrocytes convey signals that directly reshape the innervation of affected cortical brain areas and potentiate axonal communication in impaired neuronal tracts through EVs. These effects are accompanied by neurological improvement after stroke. Previous studies have shown that astrocytes are highly heterogeneous cells, with typical types accounting for ̴70% of astrocytes in culture ^64^, promoting neuron adhesion and neurite growth, while atypical astrocytes inhibit such processes ^65^. Reactive astrocytes, like those that form glial scars after brain damage, potentially limit axonal growth after stroke, mainly by producing an extrinsic inhibitory environment. Our results might also reflect the enhancement of soluble signals released by repair-promoting astrocytes, and by the external administration of these EVs, we could surpass the physical inhibitory barriers promoted by reactive gliosis.

Given the dire need for suitable biomarkers for post-stroke plasticity mechanisms, we propose that astrocyte-derived particles, which can be isolated from human plasma ^66,67^, may serve as functional markers for brain plasticity, especially in the chronic phase after stroke when neurological restoration is minimal and can only possibly be obtained by proper neurorehabilitative interventions.

In conclusion, understanding the biological basis of neurological function restoration after stroke is critical for designing intervention therapies by the exploitation of endogenously coded mechanisms for the repair of the damaged brain. The use of isolated EVs from cultured astrocytes, even if unmodified, may shorten the time requires for neurological recuperation, or even more so, extend the very limited time window of spontaneous recovery and increase the proportion of functional gains in patients. Additional studies are warranted to explore the optimal time point for astrocyte-derived EVs administrations and the optimally effective and safe way to deliver them to the damaged brain.

## METHODS

### Animals

This study used young 6-week-old (270-290 g) wild-type Wistar rats subjected to MCAO as described below. Rats were bred at the Animal Facility of IFC-UNAM certified by the Secretariat of Agriculture and Rural Development (SADER–Mexico). Animals were housed in individual cages in a 12 h light/dark cycle with food and water *ad libitum*. Rats were killed at 1, 7, 14, and 21 d post-stroke. All experimental procedures were conducted under the current Mexican law for the use and care of laboratory animals (NOM-062-ZOO-1999) with the approval of the Institutional Animal Care and Use Committee (CICUAL-IFC-LTR93-16). Experiments are reported in compliance with the Updated Animal Research: Reporting in Vivo Experiments (ARRIVE 2.0) guidelines.

### Study design

For MRI studies, animals were randomly divided into four groups (n = 4 / group). The sample size was calculated based on a previous study from our group used as pilot^63^, with *a priori* calculations to detect a medium Cohen’s d effect size > 0.3, statistical β power of 0.8 and significance of 0.05. The mortality rate was assumed to be 0.4 based on pilot experiments. These parameters were chosen to minimize the number of animals used. The inclusion criteria considered included: a reduction of blood perfusion below 50 % of basal values, which roughly corresponds to the effect of occluding the common carotid artery, reperfusion above 50 % baseline values within 5 min, total occlusion time of 60 min, absence of subarachnoid or intraparenchymal hemorrhages and survival for 21 days after stroke. For humane reasons, experiments were terminated before the indicated period if animals presented with hemiplegia or generalized weakness that made them unable to eat or drink autonomously; and such experiments were not included in the analyses. For histological evaluations at 1, 7, and 14 d, post-stroke independent experiments that met the described criteria were replicated three times. The characterization of axonal growth and construction of polar maps was performed in a single experiment per condition. The electrophysiological recordings were performed in groups of n = 4 per experimental condition. This study was limited to assess effects on male rats to limit potential confounds of estrogen-mediated neuroprotective actions present in female rodents.

### MCAO

Ischemic stroke was performed as previously reported with slight modifications^63^. Briefly, rats were subjected to isoflurane anesthesia (5 % for induction followed by ≤1.5 % during surgery) with oxygen as the carrier. Normal ventilation was autonomously maintained. A nylon monofilament with a silicone-covered tip (403734, Doccol, Sharon, MA) was inserted through the ligated left external carotid artery, and intra-luminally advanced through the internal carotid artery until it occluded the MCA. The occlusion was maintained for 60 min after which the monofilament was removed. Body temperature was maintained at 37 °C with a heating pad for the duration of surgery. At the end of the procedure, the neck’s skin was sutured, and rats were returned to their cages. During the entire experimental procedure, the cerebral blood flow (CBF) was monitored in the territory irrigated by the MCA with laser-Doppler flowmetry at the following stereotaxic coordinates; AP -1.5 L +3.5 from Bregma, with a laser-Doppler probe (model 407, Perimed, Järfälla, Sweden) connected to a Periflux System 5010 (Perimed). CBF was continuously monitored with an acquisition interval of 0.3 s using the Perisoft software (Perimed).

### Primary astrocyte cell culture

Primary cortical astrocyte cell cultures were prepared and maintained using methods similar to those described previously ^28^. Briefly, astrocytes were isolated from the cerebral cortices of postnatal day 1-2 Wistar rats. The tissue was digested with trypsin and subsequently mechanically dissociated in Hanks’ balanced salt solution. The cell suspension was plated in poly-D-lysine culture flasks containing Dulbecco’s modified Eagle’s medium/F-12 medium (Gibco BRL) and 10% fetal bovine serum (FBS) (Gibco BRL). Cultures were shaken to remove less-adherent cells, with remaining cells determined to be ̴98% GFAP^+^ astrocytes. Experiments were performed between 3 to 5 passages.

### EVs isolation and characterization

EVs were purified from astrocyte cell culture supernatants under two experimental conditions: incubated for 48 h under normoxic conditions (NxEV) or for 6 h under hypoxia, followed by 42 h for recovery in normoxia (HxEV). For hypoxia, cultures were incubated inside a hypoxic chamber (Stemcell, Stemcell Technologies Inc., Canada) with a 100% N_2_ atmosphere for 6 h at 37°C. For the collection of exosomes secreted exclusively by astrocytes, a conditioned medium was prepared with exosome-free FBS. The conditioned medium was recovered and filtered with a 220 nm pore membrane to remove large cell debris, microvesicles derived from plasma membrane with a diameter greater than 220 nm, and large apoptotic bodies. The supernatant was collected and sequentially ultracentrifuged at 50,000 x g for 30 min, followed by 100,000 x g for 70 min. The purified vesicles were resuspended in 100 µl PBS.

A fraction of the resuspended vesicles was diluted 1:50 for nanoparticle tracking analysis measurement (NTA) by Nanosight (NS300 Malvern Panalytical UK) at the National Laboratory of Flow Cytometry (IIB-UNAM, Mexico). Vesicle size was determined by estimating the diffusion coefficient calculated from the Brownian motion direct observation using the Stokes-Einstein equation and depicted as a size histogram. Both acquisition and analysis settings were kept constant for control and hypoxic samples.

For the characterization by transmission electron microscopy, 2 µL of EVs suspension were loaded onto glow-discharged 400 mesh copper/carbon-coated grids and left to settle for 5 min. After a brief wash with drops of distilled water, the grids were stained in 2% uranyl formate for 1 min and blow-dried on Whatman filter paper. Exosomes were examined with a JEOL-JEM-1200 transmission electron microscope at an accelerating voltage of 80 keV.

The expression of the exosome canonical marker CD63 was determined by Western blot. The quantification of protein content in exosome suspensions and whole-cell lysates was undertaken by a modified Lowry assay (DC Protein Assay, BIORAD). For immunoblotting, sample preparation was performed in Laemmli buffer, and 5 µg of proteins were resolved by 7.5% SDS-polyacrylamide gel electrophoresis and transferred to polyvinylidene difluoride membranes. Membranes were blocked with 5% bovine serum albumin (BSA) in Tris-buffered saline (TBS) containing 0.1% Tween 20. After blocking, the membrane was incubated overnight with the primary polyclonal antibody anti CD63 (1:200, Santa Cruz Biotechnology; sc-15363), followed by a 2 h incubation with the secondary HRP-conjugated goat anti-rabbit antibody (1:200, GeneTex; GTX213110). Following incubation with the immobilon forte substrate (EMD Millipore) the blot was scanned on a C-DiGit apparatus (LI-COR Biosciences).

### Administration of astrocyte-derived EVs in the brain

A 4 µl volume of an exosome suspension in phosphate buffer was administered by intracerebroventricular injection (i.c.v.) in the corresponding animal groups 30 min after removing the intraluminal filament in the MCAO procedure, which marked the beginning of reperfusion. The injection was performed at the following stereotaxic coordinates: AP -0.8 and L -1.5 from Bregma and V -4 from dura matter at a flux rate of 0.8 µL/min using a graduated glass microcapillary pipette pulled to produce a tip < 50 µm in diameter. The number of exosomes administered in each experiment contained equal quantities of total protein in the range of 400 ng.

For the characterization of the distribution and bioavailability of EVs injected in the brain, we stained them with PKH26 (Sigma-Aldrich), per the manufacturer’s directions, and proceeded to ICVI. We determined the presence of labeled EVs in the brain at 2 and 24 h after their administration by evaluating 40 µm paraformaldehyde (PFA)-fixed coronal brain slices. Sections were blocked and permeabilized with 0.5% TBS-T and 5% BSA and incubated overnight at 4°C with primary antibodies anti-MAP-2 (1:500, Abcam; ab32454) and anti-GFAP (1:500, Sigma; G3893), and after 3 washes, secondary Alexa fluor 647-conjugated goat anti-rabbit and Alexa Fluor 488-conjugated anti-mouse antibodies (1:500 each), and DAPI (ThermoFisher; 1:10000). The preparations were examined by confocal microscopy with an LSM 800 microscope (Zeiss, JENA, Germany) using a 63 X oil immersion objective.

### Behavioral testing

Animals were evaluated with a battery of neurological tests to assess sensorimotor deficits at 24 h, 7, 14, and 21 d after stroke. The severity of functional deficits was scored by assessing eight items described in Table 1. All evaluations were cross-validated by a trained observer blinded to the experimental treatment that analyzed the tests’ recorded videos.

### Magnetic Resonance Imaging

MRI scans were acquired at 7, 14, and 21 days post-MCAO with a 7 T MR System (Varian, Inc.), and assessments made at 24 h were performed with a 7 T MR (Pharmascan 70/16; Bruker Biospin, Ettlingen, Germany), on animals anesthetized with isoflurane (5% for induction followed by ≤2% during image acquisition). T2-weighted images were obtained with a fast spin-echo sequence; TE/TR = 35.47/3714 ms; FOV = 64 × 64 mm; matrix = 256 × 256; voxel resolution = 250 µm per side; slice thickness = 0.5 mm. T1-weighted images were acquired with a 3D gradient-echo sequence with the same orientation and resolution; TE/TR = 2.3/4 ms. Diffusion-weighted images were acquired using a spin-echo echo-planar imaging sequence: TE/TR 30.5/2500 ms; FOV = 64 × 64 mm, matrix = 64 × 64, slice thicknes = 1 mm, yielding isometric voxel resolution of 1 mm per side; diffusion gradient directions = 6, diffusion gradient duration (δ) = 5 ms, b-value = 800 s/mm^2^; an additional b = 0 s/mm^2^ image was acquires with the same parameters.

### Diffusion tensor imaging (DTI)

The diffusion tensor was fitted at each voxel for DTI analysis using DSI Studio (http://dsi-studio.labsolver.org/Manual/diffusion-mri-indices; 12/6/2015). Next, tractography was performed using a deterministic fiber-tracking algorithm ^68^. Seed regions were placed at the striatum and corpus callosum. The anisotropy threshold was 0.21, and the angular threshold was 30 degrees. The step size was 0.7 mm, averaging the propagation direction with 30 % of the previous direction smoothed the fiber trajectories. A total of reconstructed streamlines (tracts), AD, MD, and RD, were calculated by area.

### Immunofluorescence and confocal microscopy

For immunohistological analyses, three rats per group were anesthetized with pentobarbital (100 mg/kg) and transcardially perfused with 200 mL ice-cold 0.9 % NaCl followed by 250 mL ice-cold 4% PFA. Brains were collected and post-fixed in 4 % PFA for 24 h and then cryoprotected in 30 % sucrose. Whole PFA-fixed brains were cut into 40 µm thick sections in a cryostat to produce ten series of consecutive sections that were 400 µm apart. Brain sections containing infarct core and penumbra were blocked with 5% BSA in TBS with 0.5% v/v Triton X-100 (TBS-T) and incubated with anti-MAP-2 (1:200; Invitrogen, PAS-17646) and anti-TUJ-1 antibodies (1:200; Merck-Millipore, MAB380) for 48 h. Sections were washed three times with TBS followed by 2 h incubation at RT with Alexa Fluor 488-conjugated anti-mouse and Alexa 546-conjugated anti-rabbit antibodies (1:300 each; ThermoFisher Scientific, A32723, A11035) in TBS. Images were obtained in a Zeiss LSM 800 confocal microscope using a 20X objective. An average of 40 μm for each Z-stack was obtained, every 0.5 μm of optical sectioning.

### Axonal mapping

To determine the effect of EVs in neuronal projections recovered over time, we injected 2 µl of 1% w/v FITC-conjugated cholera toxin B subunit (Sigma, C1655) in rats subjected to MCAO on day 14 post-stroke and analyzed the distribution of the labeling by confocal microscopy at 21 d post-stroke. In a separate series of experiments, one rat of each group subjected to MCAO was administered with 0.5 µl of 0.02 % 1,1’-dioctadecyl-3,3,3’,3’-tetramethylindocarbocyanine perchlorate (Dil) in 0.1% DMSO in the striatal core of the infarction (AP -0.1, ML 2.0 and DV - 4.2) on day 14 after MCAO. On day 21, the animals were intracardially perfused as indicated above with 4% PFA. The cortical hemispheres were collected, and the ipsilateral hemicortex was flattened as previously described ^69^. The flattened cortices were clarified in a sucrose gradient (15, 30, 45, 60, and 75 %) with Triton X-100 (0.4, 0.6, 0.8, and 1 %) for 15 d per stage. For two-photon microscopy, the flattened cortices were immersed in 2% agarose between glass slides and scanned with a Zeiss LSM710 microscope to acquire a series of images in a stack of the whole cortex using a C-Apochromat 10X/0.45 W M27 objective with a pixel dwell of 6.3 ms at a wavelength of 860 nm.

Stack maximum intensity projections for each image were binned in a 20 × 20 square calculating the pixels’ median intensity inside each bin to create a new pixel. The images were processed for noise reduction using a scikit-image median filter ^70^, calculating the median pixel value in a 2-pixel-radius disk. We set a threshold filter at percentile 90-95 of the pixel values for establishing the positive signal intensity and kept only the pixels with the highest values. The pixel cartesian coordinates were then obtained, setting the center (0,0) at the first branch split of the M4 segment in the MCA’s superior trunk in the barrel cortex, identified by cytochrome oxidase staining (Supplementary Figure 6). This point was considered as the origin of the polar transformation. All the processing and plotting were undertaken with ad hoc Python scripts available at https://github.com/TYR-LAB-MX/PolarPlot_HerasRomero_2021.

### In vivo electrophysiological recording

Input-output (I/O) curves were assessed 21 d after stroke. For this, animals (n=4 per group) were anesthetized with isoflurane 5% for induction, followed by ≤2% during surgery. Body temperature was maintained at 35° C with heating pads. Rats were placed on a stereotaxic frame, and the skull was exposed. A constant current was delivered by direct and unilateral stimulation of the corpus callosum using a bipolar stainless-steel electrode placed at the stereotaxic coordinates AP +0.2, ML -1.0, DV -3.7 from Bregma, with a Grass S48 stimulator and a photoelectric stimulus isolation unit Grass PSIU6 (Grass Instrument Co. Quincy, MS). Corpus callosal responses were recorded unilaterally with a monopolar stainless-steel electrode (127 μm diameter) placed at the stereotaxic coordinates AP +0.2, ML +1.0, DV -3.7 from Bregma. The evoked responses were measured with the compound action potentials (CAP) amplitude, measured in negative peak 1 (N1) and peak 2 (N2). I/O curves were built with threshold folding of intensity (1-10 X) to determine the axonal conduction for a range of stimulation intensities. The threshold was defined at the stimulation intensity required to produce a 0.10 mV amplitude response in N1. The electric signal was digitalized, stored, and analyzed using the software DataWave SciWorks (Broomfield, CO, USA).

### Statistics

GraphPad Prism 8 was used to analyze all data. The normal distribution in each data set was corroborated using the Shapiro-Wilk normality test. Neurological scores and DTI parameter changes among experimental groups and over time within each group were tested on a 2-way analysis of variance ANOVA with repeated measures based on a general linear model, followed by Tukey’s post hoc test. Data were considered significant at α ≤ 0.05 level.

## Data Availability

The datasets used and/or analyzed during the current study are available from the corresponding author on reasonable request.

## Ethics Statement

All the animal experimentation was carried out at the Universidad Nacional Autónoma de México in Mexico City and Juriquilla Qro., Mexico, in accordance with the Mexican law for the use and care of laboratory animals (NOM-062-ZOO-1999) with the approval of the Institutional Animal Care and Use Committee (CICUAL-IFC-LTR93-16), and is in compliance with the guidelines for animal experimentation of the National Research Council (Committee for the Update of the Guide for the Care and Use of Laboratory Animals, 2011) and the National Institutes of Health of the United States of America (DHEW publication 85-23, revised, 1995).

## Author contributions

YHR and LTyR conceived the project and designed the experiments. YHR performed the MCAO surgeries. YHR, AM, and LC performed MRI assessments, and AM produced the tractography. RSM made two-photon sample preparation and acquisition, APH and RRH analyzed the images and produced polar plots. IP carried out the proteomic meta-analysis. YHR and RRH made histological sample preparations and image acquisition. ACR performed the exosome characterization, and BNBV analyzed the distribution of EVs in the brain. Electrophysiological experiments were performed by YHR, AMM, EU, and MLE. YHR, AM, RRH, APH, PMC, NHG, MLE, LC, and LTyR analyzed the data. LTyR wrote the manuscript with input from YHR, PMC, NHG, MLE, and LC. All authors read and approved the final version of the manuscript.

## Acknowledgments

We thank Cristina Aranda Fraustro and Dr. Alfredo Cárdenas for their assistance with experimental procedures, Dr. Alicia Güemez-Gamboa (Northwestern University) and Elliot Glotfelty (NIA/NIH, Karolinska Universitat) for critically reviewing the manuscript, Dr. Abraham Rosas-Arellano, for assistance with the acquisition of confocal images and post-processing, Dr. Yazmín Ramiro-Cortés and Gerardo Perera for their expert assistance with two-photon image acquisition, Claudia Rivera-Cerecedo for animal care, Daniela Rodríguez-Montaño for assistance with histological sample preparations, the National Laboratory for Magnetic Resonance Imaging, particularly Dr. Juan Ortiz-Retana for technical assistance, and Dr. Gloria Soldevila and Dr. Cynthia López Pacheco of the Laboratorio Nacional de Citometría de Flujo (IIB-UNAM) for their helpful assistance with NTA. The corresponding author is deeply grateful to Pilar Martínez, Mario Arredondo, Claudia Islas, Pablo Montiel, and Angel Cedillo for their outstanding assistance with the administrative burden that kept the laboratory running during the difficulties that arose through the COVID19 pandemic shutdown. Yessica Heras-Romero conducted this study in partial fulfillment of the requirements of the Programa de Doctorado en Ciencias Biomédicas of Universidad Nacional Autónoma de México.

---

**Supplementary figure 1.**
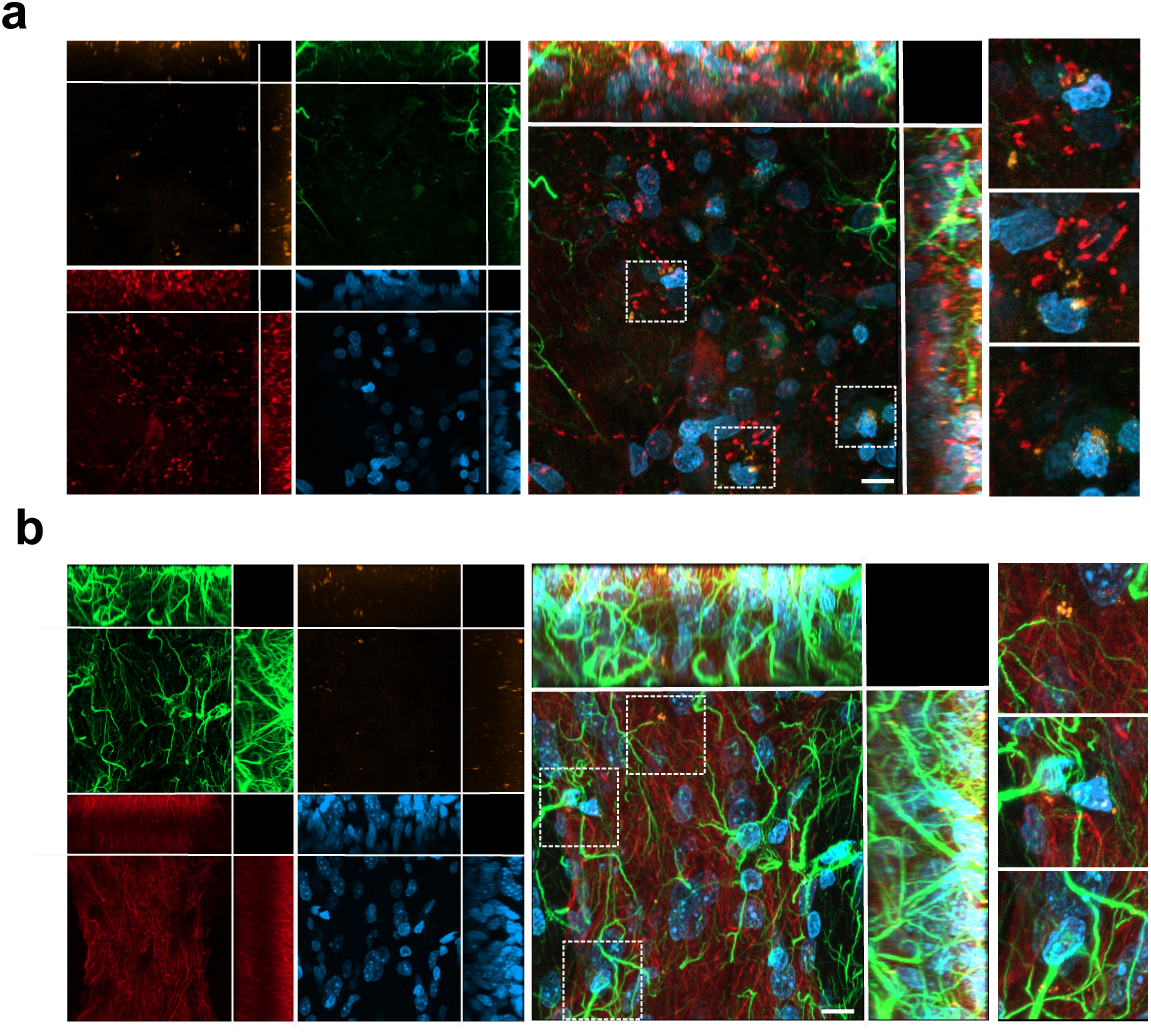
Distribution of EVs stained with PKH26 (orange) at 24 h after i.c.v. injection in the brain of stroke-challenged rats in the striatum **(a)** and motor cortex **(b)**. EVs internalize in neurons (MAP2; red) and astrocytes (GFAP; green) and preferentially localized to perinuclear (DAPI; blue) regions. Dotted squares demark regions where EVs localize. Images are maximum projections of a Z-stack of 20 optical slices showing the orthogonal planes, and nuclei are stained with DAPI (blue), scale bar equals 10 µm.

**Supplementary figure 2.**
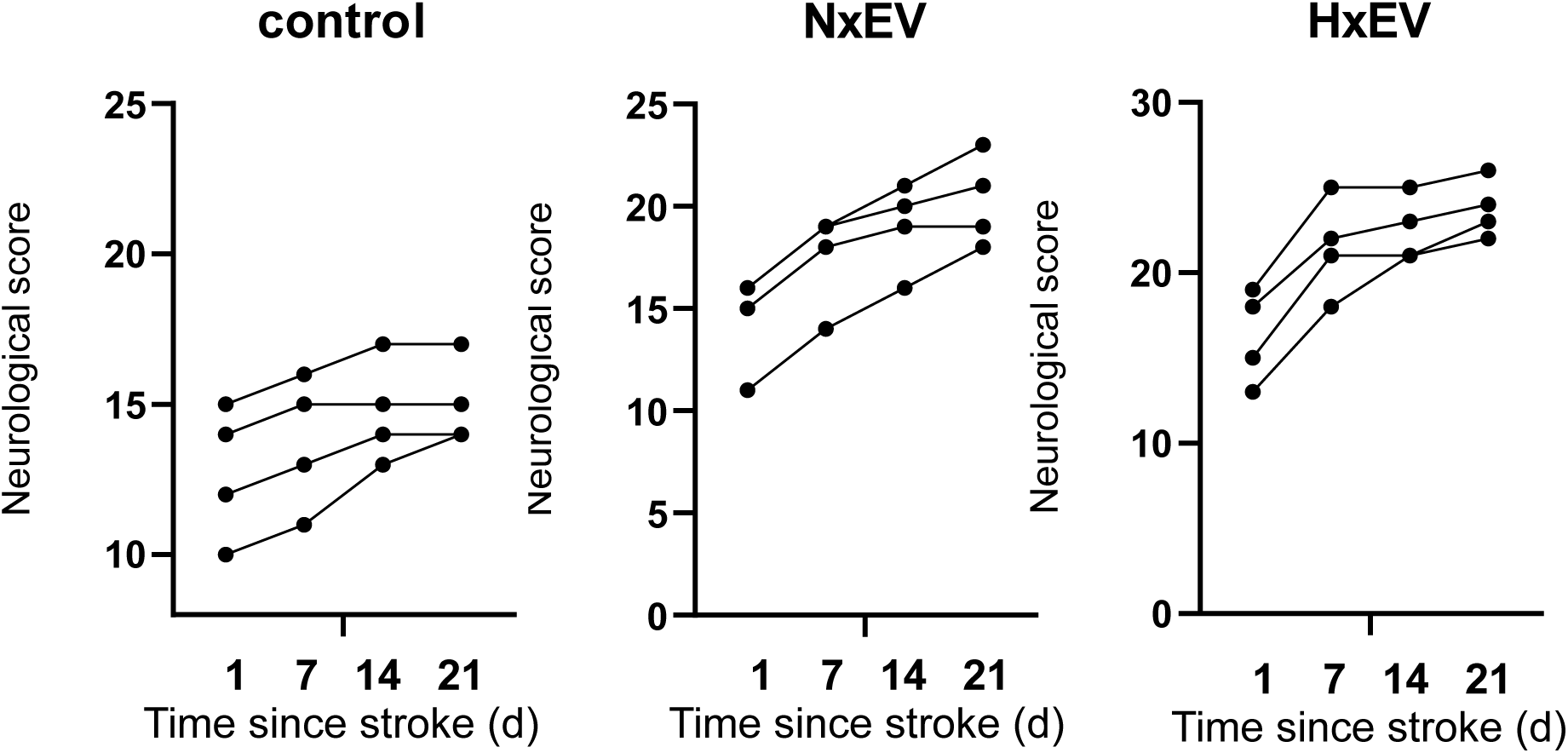
Spontaneous recovery trends of individual animals in each group, assessed at 1, 7, 14, and 21 days post-stroke.

**Supplementary figure 3.**
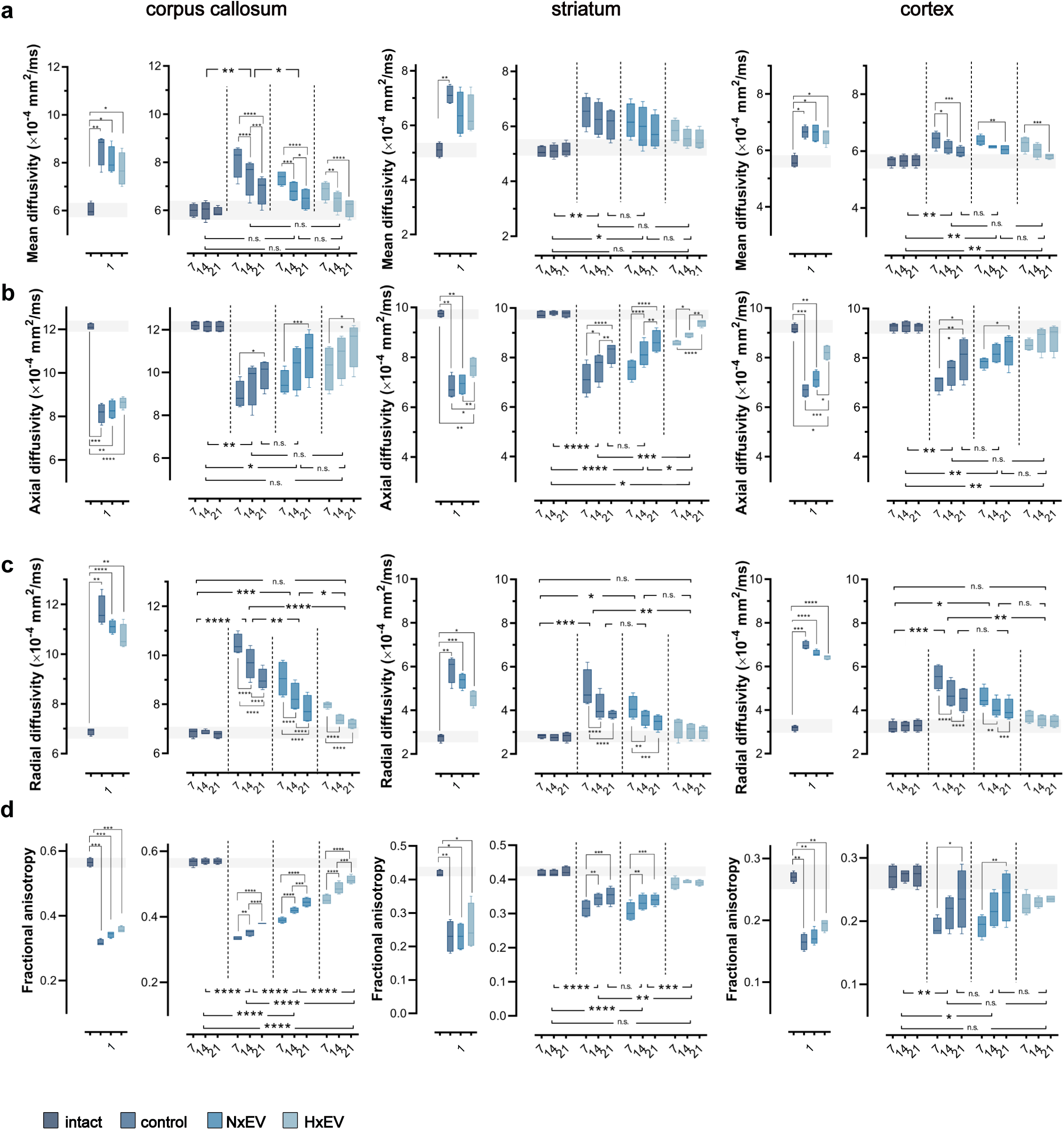
Evolution of DTI parameters in the contra-lesional hemisphere. **(a)** mean diffusivity, **(b)** axial diffusivity, **(c)** radial diffusivity, and **(d)** fractional anisotropy were determined from diffusion tensor imaging (DTI) of the contralateral corpus callosum (left column), striatum (middle column), and motor cortex (right column) at 1, 7, 14 and 21 days post-stroke. Boxplots on day 1 show the alterations caused by the stroke in all four DTI parameters; no statistical differences exist between stroke-challenged animals treated with vehicle (control) and those that received EVs 30 min after the initiation of reperfusion. Boxplots show the min and max values within each group, the dispersion span from Q1 to Q3, and the mean, n=4. The shaded horizontal bar in each plot marks the span of ± 1 S.D. of the intact group baseline values. Statistical differences of the recovery trend are indicated among groups with repeated measures two-way ANOVA followed by Tukey’s post hoc, and changes over time within each group are also indicated with two-way ANOVA followed by Tukey’s post hoc. * p<0.05, **p<0.01, ***p<0.001 and ****p<0.0001.

**Supplementary figure 4.**
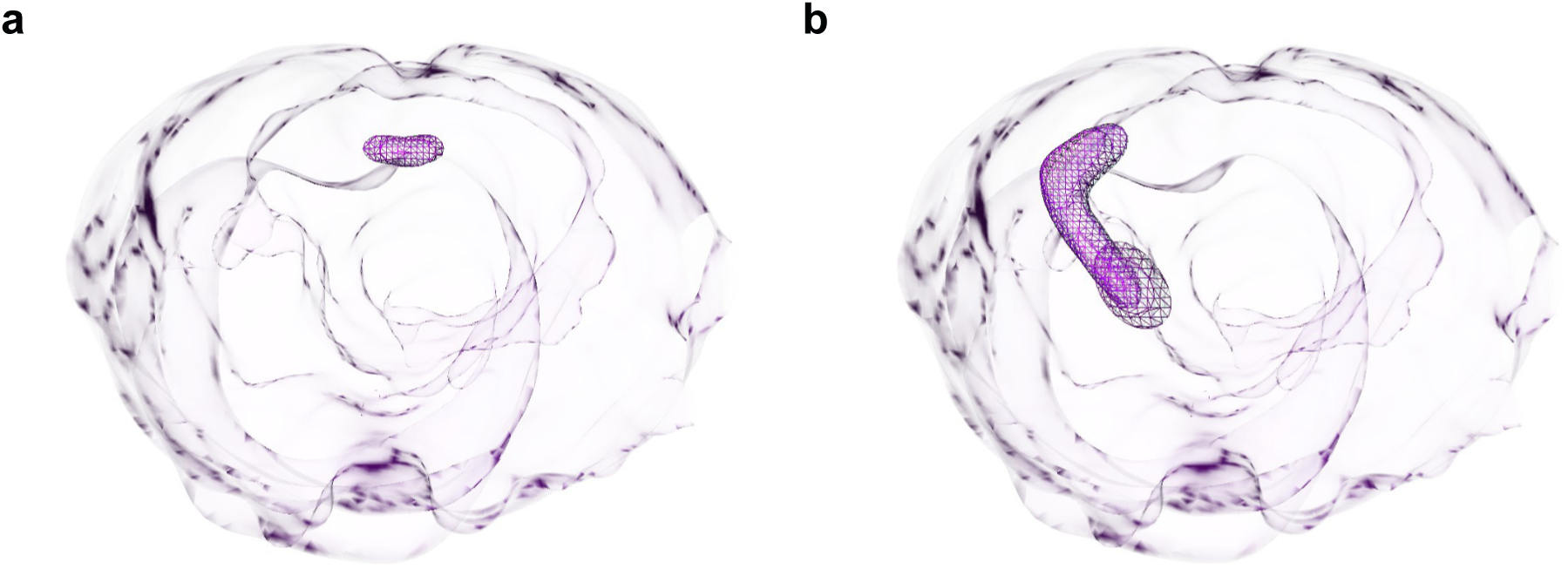
Seed regions used for tractography of the corpus callosum **(a)** and the corticostriatal **(b)** tracts.

**Supplementary figure 5.**
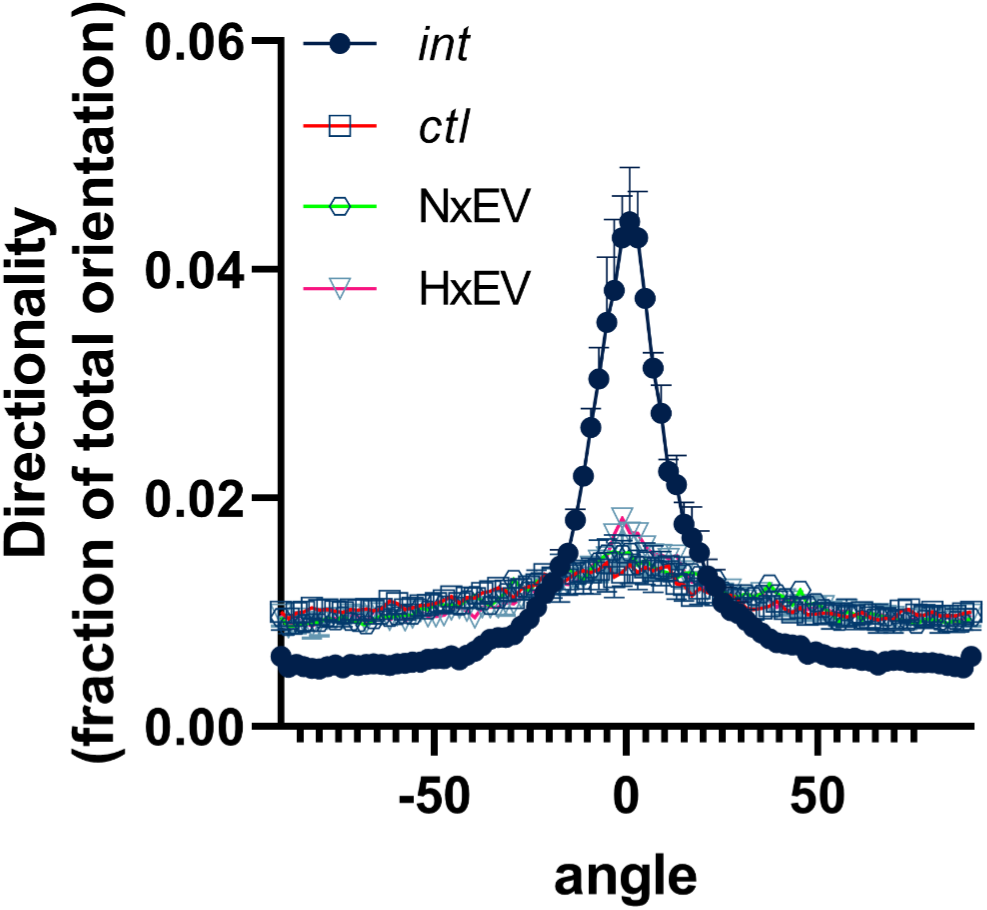
Analysis of the directionality of MAP-2 immunolabeled fibers in the striatum 21 days after stroke. The EV-induced increases in fiber staining did not restore the directionality of the fibers after stroke.

**Supplementary figure 6.**
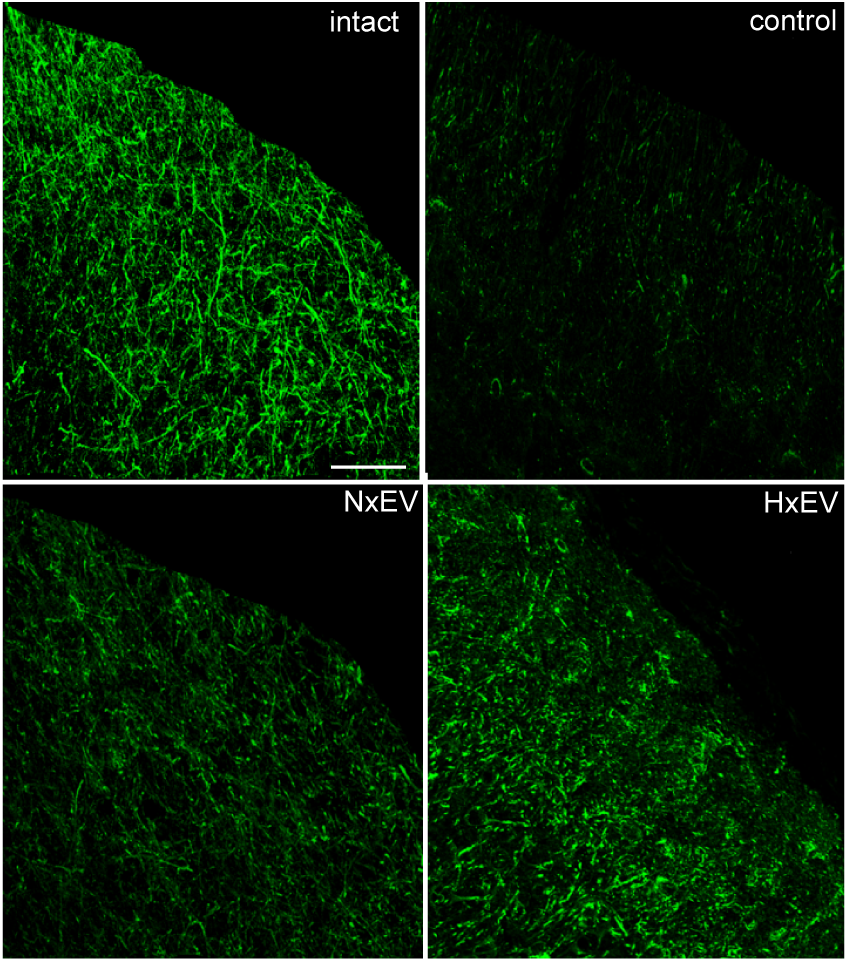
Representative confocal micrographs of cortical labeling in animals injected in the dorsal striatum with the cholera toxin subunit B, which is retrogradely transported through the axonal system. Injections were made on day 14 after stroke and followed the fluorescent label’s localization in the innervated cortical regions at day 21 post-stroke. Scale bar equals 50 µm.

**Supplementary figure 7.**
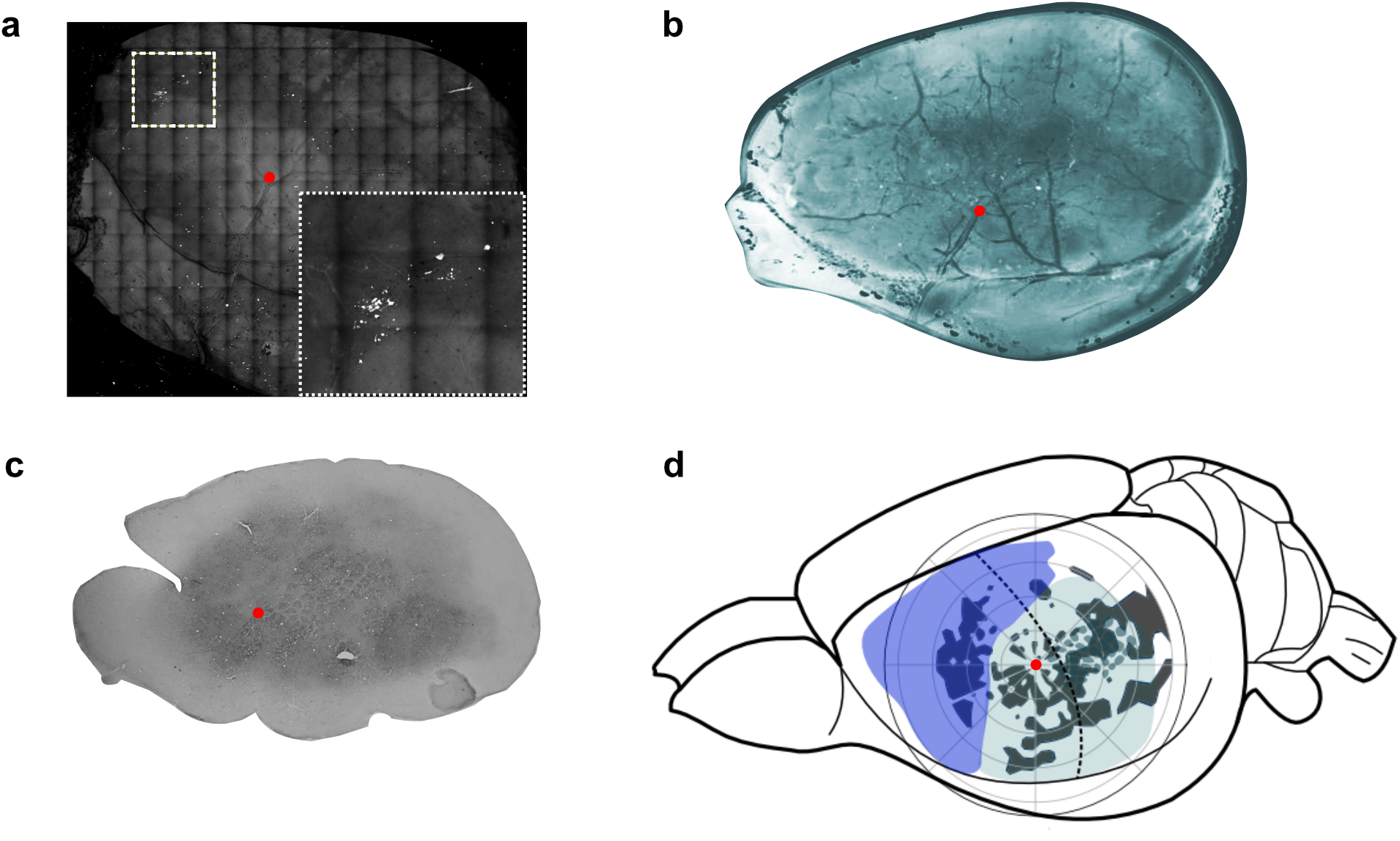
Mapping of the cortical axonal projections from the dorsal striatum was made by superposing the panoramic two-photon image **(a)** where the vasculature can be clearly appreciated, with the ink-stained flattened cortex showing the superficial cortical vasculature **(b)** and the cytochrome c oxidase staining of the barrel cortex **(c)**. The red dot indicates the exact anatomical localization at the first branch split of the M4 segment in the MCA’s superior trunk and its relative position to the barrel cortex. This anatomical location was designated as the origin (0,0 coordinate) for polar plots **(d)**.

